# Native doublet microtubules from *Tetrahymena thermophila* reveal the importance of outer junction proteins

**DOI:** 10.1101/2022.09.30.510376

**Authors:** Shintaroh Kubo, Corbin Black, Ewa Joachimiak, Shun Kai Yang, Thibault Legal, Katya Peri, Ahmad Khalifa, Avrin Ghanaeian, Melissa Valente-Paterno, Chelsea De Bellis, Phuong Huynh, Zhe Fan, Dorota Wloga, Khanh Huy Bui

**Affiliations:** Department of Anatomy and Cell Biology, Faculty of Medicine and Health Sciences McGill University, Québec, Canada; Laboratory of Cytoskeleton and Cilia Biology, Nencki Institute of Experimental Biology of Polish Academy of Sciences, 3 Pasteur Str, 02-093 Warsaw, Poland

**Author notes:** These authors contributed equally to this work (Shintaroh Kubo, Corbin Black, Ewa Joachimiak).

## Abstract

Cilia are ubiquitous eukaryotic organelles responsible for cellular motility and sensory functions. The ciliary axoneme is a microtubule-based cytoskeleton consisting of two central singlets and nine outer doublet microtubules. Cryo-electron microscopy-based studies have revealed a complex network inside the lumen of both tubules composed of microtubule-inner proteins (MIPs). However, the functions of most MIPs remain unknown. Here, we present single-particle cryo-EM-based analyses of the *Tetrahymena thermophila* native doublet microtubule and identify 38 MIPs. These data shed light on the evolutionarily conserved and diversified roles of MIPs. In addition, we identified MIPs potentially responsible for the assembly and stability of the doublet outer junction. Knockout of the evolutionarily conserved outer junction component CFAP77 moderately diminishes *Tetrahymena* swimming speed and beat frequency, indicating the important role of CFAP77 and outer junction stability in cilia beating generation and/or regulation.

## Introduction

Cilia are hair-like structures protruding from the cell surface and are conserved from protists to humans. Immotile sensory cilia play an essential role in sensing and transducing external signals such as sonic hedgehog from the surrounding environment to the cell. Coordinated beating of motile cilia enables transport of cells or fluids along the surface of ciliated cells, and thus are key factors in clearing mucus out of the respiratory tract or circulating cerebrospinal fluid (Reiter & Leroux, 2017). The core structure of motile cilia, the axoneme, consists of two central singlet microtubules and nine peripherally positioned doublet microtubules (DMT) (Bui, Sakakibara, Movassagh, Oiwa, & Ishikawa, 2008; Nicastro et al., 2006). The assembly and stability of the DMTs is indispensable for the ciliary function as they serve as a scaffold for numerous ciliary complexes, the force transducers for ciliary bending (Muneyoshi Ichikawa et al., 2019), and tracks for intraflagellar transport (Stepanek & Pigino, 2016).

The inner (luminal) wall of the DMT is supported by an intricate network of microtubule inner proteins (MIPs) (M. Ichikawa et al., 2017; Maheshwari et al., 2015; Nicastro et al., 2011). Using single-particle cryo-electron microscopy (cryo-EM), the identities of over 30 MIPs have been revealed in the ciliate *Tetrahymena thermophila* (Muneyoshi Ichikawa et al., 2019; Khalifa et al., 2020; S. Li, Fernandez, Fabritius, Agard, & Winey, 2022), the green algae, *Chlamydomonas reinhardtii* (Ma et al., 2019), and bovine respiratory cilia (Gui et al., 2021). Interestingly, some MIPs also contact the DMT outer surface and additional axonemal components; mutations in FAP127/MNS1 result in defects in outer dynein arm docking complex assembly (Ta-Shma et al., 2018).

At least half of the MIPs identified in those studies are conserved, including the inner junction proteins FAP20, PACRG (Dymek et al., 2019; Khalifa et al., 2020; Yanagisawa et al., 2014), FAP45, FAP52 (Owa et al., 2019), and the protofilament (PF) ribbon proteins, RIB43a and RIB72 (Ikeda et al., 2003; Norrander, deCathelineau, Brown, Porter, & Linck, 2000). Interestingly, some MIPs show poor conservation. For example, the tektin bundles of the PF ribbon were identified in bovine respiratory cilia, but not in *Tetrahymena* or *Chlamydomonas* (Gui et al., 2021).

There are few studies exploring the functions of MIPs. RIB72/EFHC1, an EF-hand protein, is implicated in juvenile epilepsy disorder (King, 2006; Suzuki, Inoue, & Yamakawa, 2020) and is important for *Tetrahymena* cell motility (Fabritius et al., 2021; Stoddard et al., 2018). FAP45 and FAP52 play a role in anchoring the B-tubule to the A-tubule wall; a lack of FAP45/FAP52 leads to B-tubule instability (Owa et al., 2019). Pierce1 and 2, the orthologs of *Chlamydomonas* FAP182 (Ma et al., 2019), are important for outer dynein arm assembly in mammals and zebrafish (Gui et al., 2021).

Until now, the identity of MIPs at the outer junction of the DMT remains unknown. The outer junction is thought to be the site of B-tubule assembly (Schmidt-Cernohorska et al., 2019). Disruption of the outer junction leads to DMT damage and consequently dysfunctional cilia. At the outer junction, α- and β-tubulins from PF B1 form a non-canonical interaction with tubulins of A-tubules (M. Ichikawa et al., 2017). Cleavage of the C-terminal tails (CTTs) of tubulins by subtilisin enables B-tubule formation *in vitro* (Schmidt-Cernohorska et al., 2019). On the other hand, CTTs of tubulin are essential for ciliary function *in vivo* (Bosch Grau et al., 2013; Vent et al., 2005). Therefore, there must be a mechanism that suppresses the C-terminal tails of tubulins of the A-tubule enabling the formation of the B-tubule.

We hypothesized that MIPs belong to one of two groups: evolutionarily conserved “core” MIPs, which function in the assembly of cilia; or species or lineages-specific “auxiliary” MIPs that add stability to the DMT. Here, we used an integrated approach combining single-particle cryo-EM, mass spectrometry, and artificial intelligence to model MIPs in the DMT lumen of the ciliate *Tetrahymena thermophila*. We showed that nearly half of the MIPs identified in *Tetrahymena* are evolutionarily conserved. In contrast to DMTs in *Chlamydomonas reinhardtii* flagella (Ma et al., 2019) and bovine respiratory cilia (Gui et al., 2021) where the MIP network adopts a 48-nm periodicity, the *Tetrahymena* MIP network repeats every 96-nm. In addition, we observed filaments on the outer surface of the DMT, which have implications for intraflagellar transport. Finally, using a combination of genetics and microscopy techniques, we revealed that CFAP77, the conserved MIP found at the outer junction of the DMT, is important for DMT stability and hence, ciliary motility.

## Results

### *Tetrahymena* DMT consists of conserved and non-conserved MIPs

We optimized the purification of DMTs from isolated *Tetrahymena* cilia to preserve all the associated structures (Materials & Methods). This approach was successfully used prior to obtain the outer dynein arm-bound DMT structure (Kubo et al., 2021). The single-particle cryo-EM analyses of the DMT from wild-type cells allowed us to obtain a structure with a global resolution of 4.1 Å (Fig. 1 and Supplementary Fig. 1). The application of focus refinement of different selected regions of the DMT resulted in maps with a resolution range from 3.6 to 3.9 Å (Materials and Methods).

**Figure 1.**
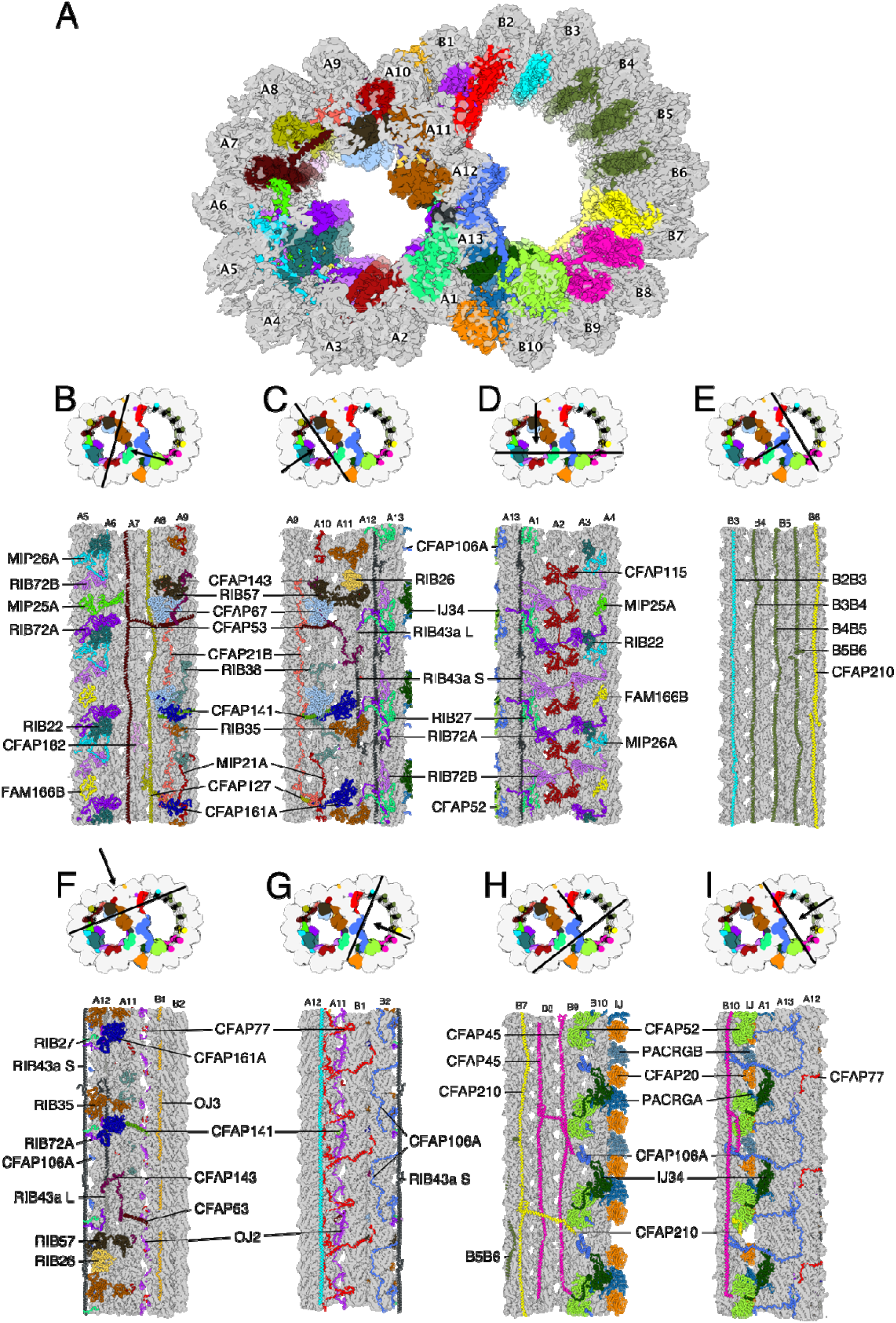
The structure of the native DMT from *Tetrahymena thermophila*. (A) A cross-section of the DMT map. Each color denotes an individual MIP. Tubulin is in gray. (B-I) Views of the lumen of the DMT from different angles as indicated by the black arrow. The cutting plane indicated by black lines.

To identify and model the MIP network, we devised a strategy that employed (i) homology modelling of conserved MIPs and (ii) de novo identification and modelling of unknown MIPs using artificial intelligent backbone tracing DeepTracer (Pfab, Phan, & Si, 2021) of densities in our map, artificial intelligent structure prediction ColabFold (Jumper et al., 2021; Mirdita, Ovchinnikov, & Steinegger, 2021) of all proteins present in the proteome of the DMT, and structure similarity search of protein backbones within the predicted structures of the proteome of the DMT (Materials & Methods, Supplementary Fig. 1). With such an approach, we could identify almost all *Tetrahymena* MIPs and show that they are forming a weaving network inside the DMT. Many of those MIP densities were missing in our previously reported structure of salt-treated DMT (Muneyoshi Ichikawa et al., 2019) such as TtPACRGA/B, TtCFAP20, TtOJ3, TtRIB22, TtRIB27.

Among the 38 identified proteins (Fig. 1, Supplementary Fig. 1, Table S1 and S2, Supplementary Movie 1), approximately half are conserved in different species (the “core” MIPs) while the remaining are non-conserved proteins (the “auxiliary” MIPs). Interestingly, several MIPs represented by a single ortholog in *Chlamydomonas* and bovine, had two paralogs, A and B, in *Tetrahymena* (Table S1). For example, TtRIB72, TtPACRG, TtCFAP106, TtCFAP161, TtCFAP21 and TtCFAP77 (see Table S1 and S2). In a few cases such as TtCFAP106 and TtCFAP77, both paralogs were detected in cilia by mass spectrometry, but we could not distinguish their densities using cryo-EM. It is possible that both paralogs co-localize along the entire cilium length or that paralogs differentially localize to proximal or distal ends. Therefore, we modelled only one of each paralog (Table S1).

FAP52, FAP45, FAP106, PACRG, and FAP20 (Fig. 2A) are core MIPs of the inner junction region, in both *Tetrahymena* and *Chlamydomonas* (Khalifa et al., 2020; Ma et al., 2019), and bovine respiratory cilia (Gui et al., 2021). Interestingly, in *Tetrahymena*, CFAP52 is stabilized by an ancillary protein, IJ34 (this study) while in *Chlamydomonas* and bovine respiratory cilia FAP52 is stabilized by auxiliary MIP FAP276 (Khalifa et al., 2020; Ma et al., 2019) and EFCAB6 (Gui et al., 2021) respectively. The two *Tetrahymena* paralogs of PACRG, TtPACRGA and TtPACRGB differ in their N-terminal region and are positioned alternately in the inner junction. In TtPACRGA, the N-terminal fragment forms helix bundles with the main HEAT fold and potentially interacts with TtCFAP20 and tubulins at the inner junction (Supplementary Fig. 2). In contrast, TtPACRGB lacks the N-terminal fragment. Consequently, the *Tetrahymena* inner junction has a 16-nm periodicity, which clearly defines the bimodality of the interdimer distance first reported in *Tetrahymena* (Muneyoshi Ichikawa et al., 2019). The lack of the N-terminal region in PACRGB might explain why the inner junction of *Tetrahymena* is less stable upon salt treatment compared to *Chlamydomonas* (Khalifa et al., 2020), pointing to the role of the N-terminal fragment of PACRGA in the inner junction stabilization. We did not observe the unknown MIP densities observed from the subtomogram averaged map of *Tetrahymena* axonemal doublet (S. Li et al., 2022) (Fig. 1A). Perhaps, it is due to the partial decoration of this protein.

**Figure 2.**
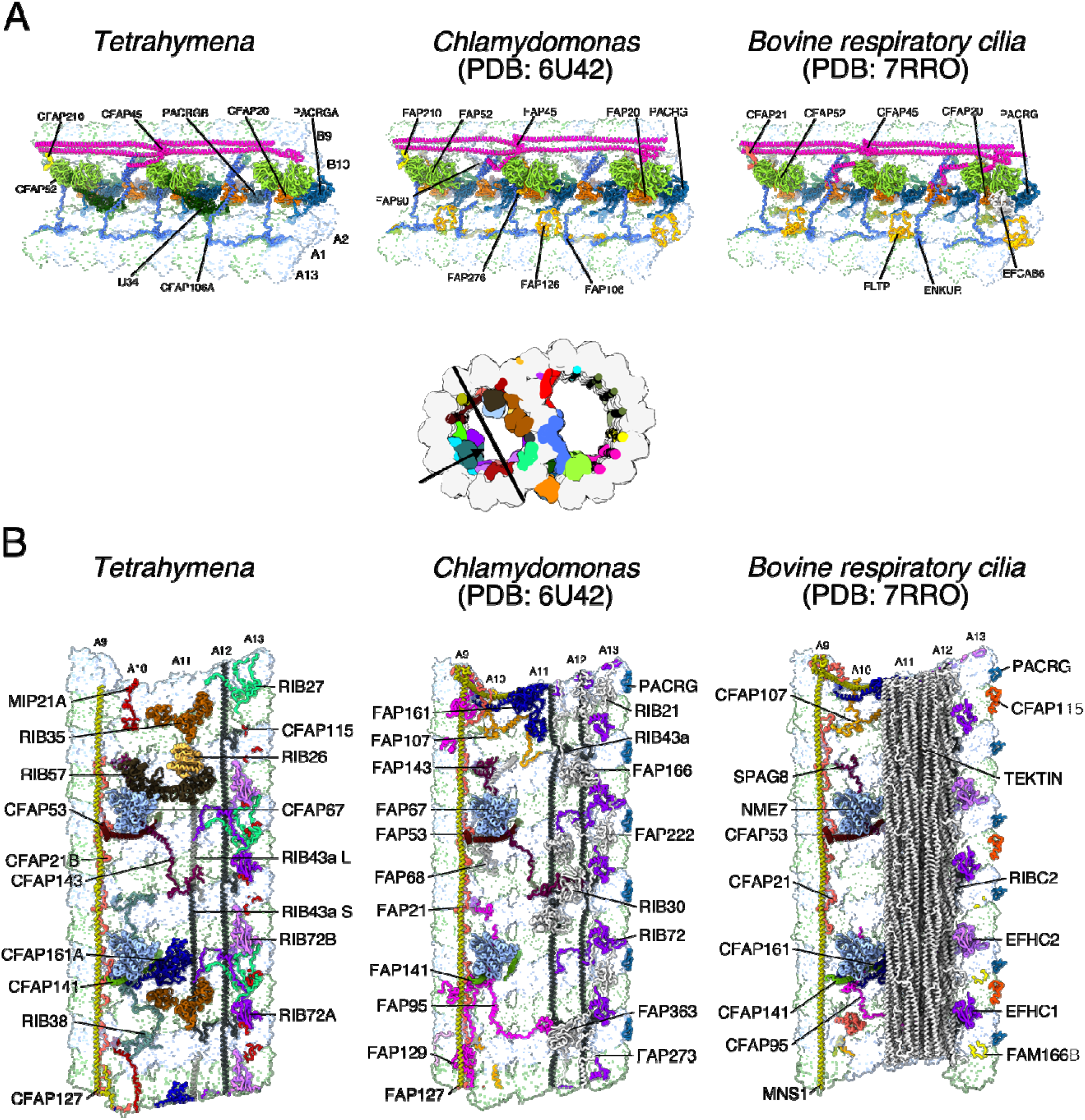
Comparison of the DMT structure from *Tetrahymena, Chlamydomonas* and bovine respiratory cilia. (A) The inner junction, note that the architecture is well conserved. (B) The PF-ribbon region, note many species-specific MIPs.

The PF ribbon is the most divergent region in the DMT (Fig. 2B). RIB43a, the core filament MIP, is conserved across species and was reported previously to be crucial for the stability of the PF ribbon region (Muneyoshi Ichikawa et al., 2019; Norrander et al., 2000). In *Tetrahymena*, the PF ribbon region is composed also of TtRIB72A, TtRIB72B, TtCFAP143, TtRIB27, TtRIB57, TtRIB35, and TtRIB26 (Fig. 2B). However, half of those proteins, TtRIB27, TtRIB57, TtRIB35, and TtRIB26 are not conserved in *Chlamydomonas* (Ma et al., 2019) and bovine respiratory cilia (Gui et al., 2021) (Supplementary Table 1). Moreover, in bovine respiratory cilia, the PF ribbon is accompanied by the tektin bundles (Fig. 2) (Gui et al., 2021). The divergence in the PF ribbon components between species suggests that the stability of the PF region is achieved through different set of MIPs instead of having strict conservation of proteins.

We also observed similarities and differences in the *Tetrahymena*, *Chlamydomonas*, and bovine respiratory cilia outer junction structure and protein composition (discussed later). To summarize, each DMT region has core components while other MIPs act as auxiliary members, possibly to reinforce the function of the core component.

### MIP distribution in *Tetrahymena* has a 96-nm periodicity

The DMT outer surface has a 96-nm periodicity regulated by the molecular ruler composed of CCDC39 and CCDC40 (Oda, Yanagisawa, Kamiya, & Kikkawa, 2014). In *Chlamydomonas* (Ma et al., 2019) and bovine respiratory cilia (Gui et al., 2021), the MIPs show a 48-nm periodicity. Interestingly, based on our observations, the *Tetrahymena* DMT has a 96-nm luminal repeat.

The *Tetrahymena* auxiliary MIP, TtCFAP115, is almost four times the size of the CrFAP115 (110kD vs. 26kD). ColabFold (Mirdita et al., 2021) prediction of TtCFAP115 shows that it contains four EF-hand pair domains (Interpro IPR011992) while CrFAP115 has only one EF-hand pair domain and repeats every 8-nm unit. We expected TtCFAP115 should have a 32-nm periodicity. To accurately position TtCFAP115, we modelled it in a 96-nm map combining several datasets that were obtained at 3.7 Å resolution for *Tetrahymena* wild-type CU428 strain, and K40 acetylation-deficient strains with either α-tubulin point mutation *K40R* (Gaertig et al., 1995; Yang et al., 2022) or knocked out tubulin acetyltransferase *MEC-17* (Akella et al., 2010). As expected, TtCFAP115 shows a clear 32-nm repeat (Supplementary Fig. 3). Since TtCFAP115 repeats with 32-nm, other MIPs such as TtRIB43a repeat with 48-nm, and TtCFAP45 repeats with 96-nm, the *Tetrahymena* DMT has a 96-nm luminal repeat consisting of three TtCFAP115 molecules per periodicity (Fig. 3A, B).

**Figure 3.**
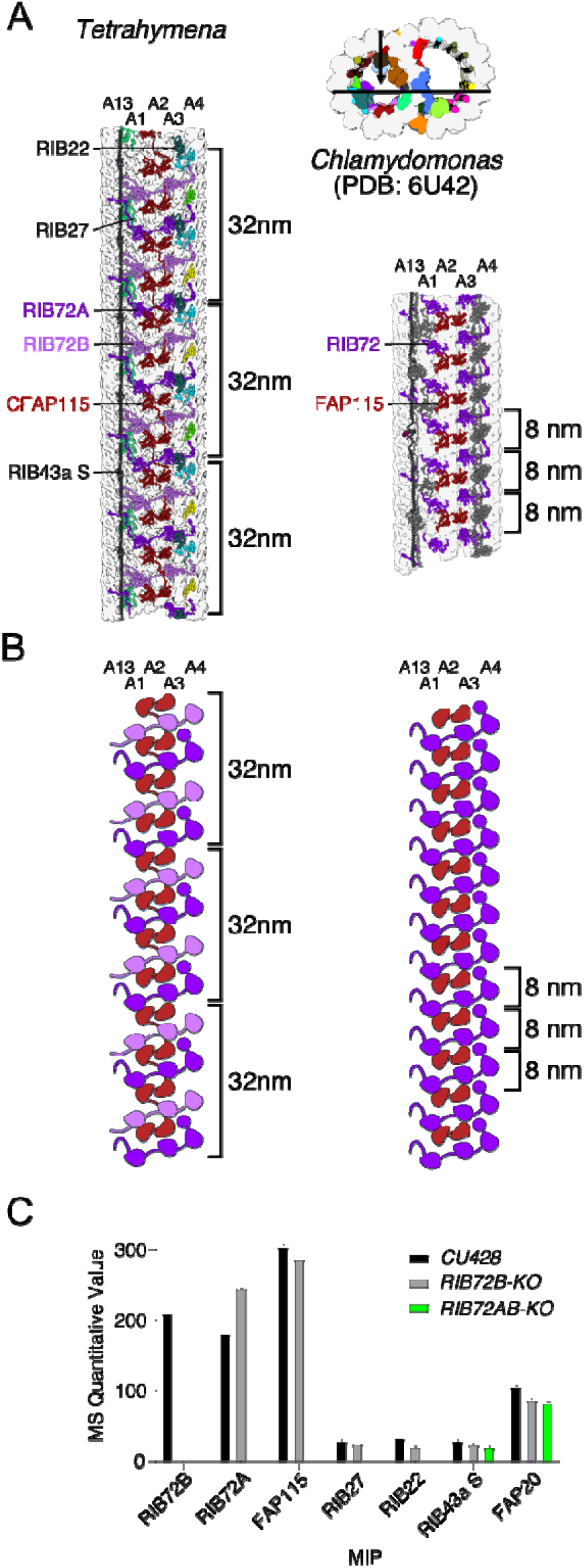
MIPs in *Tetrahymena* exhibit a 96-nm periodicity. (A) TtCFAP115 shows 32-nm repeats leading to the true 96-nm periodicity of the MIPs of *Tetrahymena*. The FAP115 in *Chlamydomonas* only repeats with 8-nm periodicity, leading to the normal 48-nm repeat. (B) The cartoon of *Tetrahymena* and *Chlamydomonas* PFs A13-A4 showing the arrangement of FAP115 and RIB72. (C) Quantitative value of MS (normal total spectra value) of WT, *RIB72B-KO* and *RIB72A/B-KO* cilia showing that TtCFAP115 is still intact after the knockout of *RIB72B*.

A single TtCFAP115 interacts with two TtRIB72A and two TtRIB72B molecules (Fig. 3A and Supplementary Fig. 3). To support our model predicting the interaction between TtRIB72 and TtCFAP115, we analysed ciliary proteomes of *RIB72B-KO* and *RIB72A/B-KO Tetrahymena* knockout mutants (Stoddard et al., 2018) by mass spectrometry (Fig. 3C and Table S3). Interestingly, TtCFAP115 was only missing in *RIB72A/B* double knockout mutant cilia suggesting that TtRIB72A could be sufficient to stabilize TtCFAP115. Our proteomic analysis of cilia of *RIB72B-KO* and *RIB72A/B-KO* mutants showed that TtMIP26A, FAM166B, TtRIB22 and TtRIB27 are also only missing in the *RIB72A/B* double knockout mutant. Thus, similar to TtFAP115, stable docking of TtRIB22 and TtRIB27 depends on TtRIB72A but not TtRIB72B (Fig. 3 and Supplementary Fig. 3).

Currently no cryo-EM maps of the DMT have shown interaction(s) that explain how the 48-nm repeat of MIPs and the 96-nm repeat of the outside proteins are established by the molecular ruler. Together with existing studies on MIPs (Ma et al., 2019) and the molecular rulers (Oda et al., 2014), our findings of the 96-nm repeat organization of *Tetrahymena* MIPs suggest that the periodicity of the inside and outside proteins/complexes can be self-regulated, probably through the head-to-tail interactions of the MIPs. Our finding showing that TtCFAP115 interacts with TtRIB72A/B is echoed by recent proteomics and low-resolution tomography studies (Fabritius et al., 2021; S. Li et al., 2022). Moreover, these authors showed that the knockout of TtCFAP115 leads to prolonged power stroke and hence decreased swimming speed, further illustrating the important roles of TtCFAP115 in *Tetrahymena* cilia, probably through DMT stabilization. Our structural analysis of TtCFAP115 highlights the importance of high-resolution reconstructions and the power of artificial intelligence prediction in the accurate validation of protein interactions, such as in the cases of missing TtRIB22 and TtRIB27 in *TtRIB72A/B-KO* (Fig. 3 and Supplementary Fig. 3). Mass spectrometry studies often reveal a broad list of missing proteins, many of which are unspecific.

### The outer surface of the intact DMT is associated with numerous MAP7-like proteins

We observed numerous filamentous structures binding to the outer wedge between two PFs with 24-nm or 48-nm repeat on the native DMT outer surface, associated with the PFs B1 to B5, and A8 to A10 (Fig. 4A, B). These PFs were shown to serve as tracks for anterograde and retrograde intraflagellar transports (Supplementary Fig. 4A).

**Figure 4.**
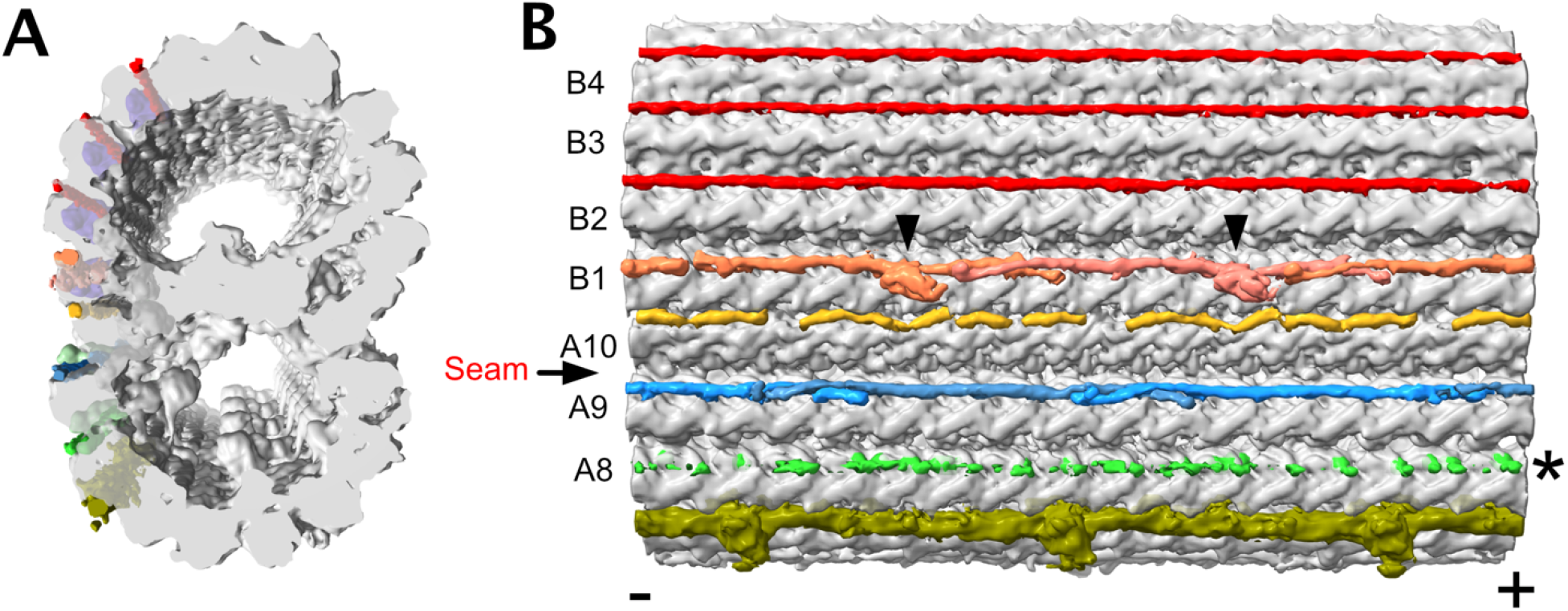
The outer surface filaments on the native DMT. (A, B) Outer surface filaments. (A) the cross section and (B) longitudinal views. The filaments bound to all pair of PFs from A8 to A10 to B1 to B5. The filament between A8A9 has lower density indicating a partial decoration. The outer surface filaments between PFs B2-B3, B3-B4, and B4-B5 appear similar and have a 48-nm periodicity. In contrast, the outer surface filaments between PFs A9-A10, A10-B1, and B1-B2 appear to have a 24-nm periodicity. These filaments have a clear head-to-tail periodic arrangement between PFs A9-A10 and B1-B2. The outer surface filament between PFs B1-B2 has a globular domain (black arrowheads). The density of the outer surface filament between PFs A8-A9 is very weak, probably due to partial decoration. (-) and (+) signs indicate the minus and plus ends of the microtubule respectively.

The low local resolution of the outer surface filament suggests that they are either partially decorated or inherently flexible. While we could not identify most of the outer surface, we were able to trace the backbone of the filament between A10 and B1 (OJ3) since it binds tightly to the outer junction wedge and thus, has better resolution. In bovine respiratory cilia and *Chlamydomonas* cilia, a similar density to OJ3 is observed but the identity of the protein is still unknown.

The outer surface filaments bind to the DMT in a similar fashion to MAP7 (Ferro et al., 2022). MAP7 is known to activate kinesin-1 even though its α-helical microtubule-binding domain recognises a site on microtubules that partly overlap with the Kinesin-1 binding site. In contrast, dynein motor activity is not impacted by MAP7 binding to microtubules (Monroy et al., 2020). In *Tetrahymena*, both intraflagellar transport motors, kinesin-2 and dynein-2, must walk on the DMT decorated by these outer surface filaments. When we overlapped the kinesin and dynein microtubule-binding domain on our DMT structure, we noticed a clear steric clash (Supplementary Fig. 4B-D), similar to the case of MAP7 (Ferro et al., 2022). Thus, likely there is a conformational change upon molecular motor binding to facilitate intraflagellar transport in the cilia.

### CFAP77 stabilizes the outer junction

Among the identified MIPs, there are proteins that support the outer junction, which is believed to initiate B-tubule formation. We observed three outer junction proteins: TtCFAP77 and OJ2 inside the B-tubule repeating every 16-nm, and OJ3 repeating with 24-nm in the wedge between A10 and B1 (Fig. 4, Fig. 5A, B, Supplementary Movie 1). Bioinformatics analyses and examination of cryo-EM maps from *Chlamydomonas* (EMDB-20631) and bovine respiratory cilia (EDMB-24664) showed that OJ2 is *Tetrahymena-specific*. In contrast, CFAP77 is conserved in many species including humans and *Chlamydomonas* (Supplementary Fig. 5A, B).

**Figure 5.**
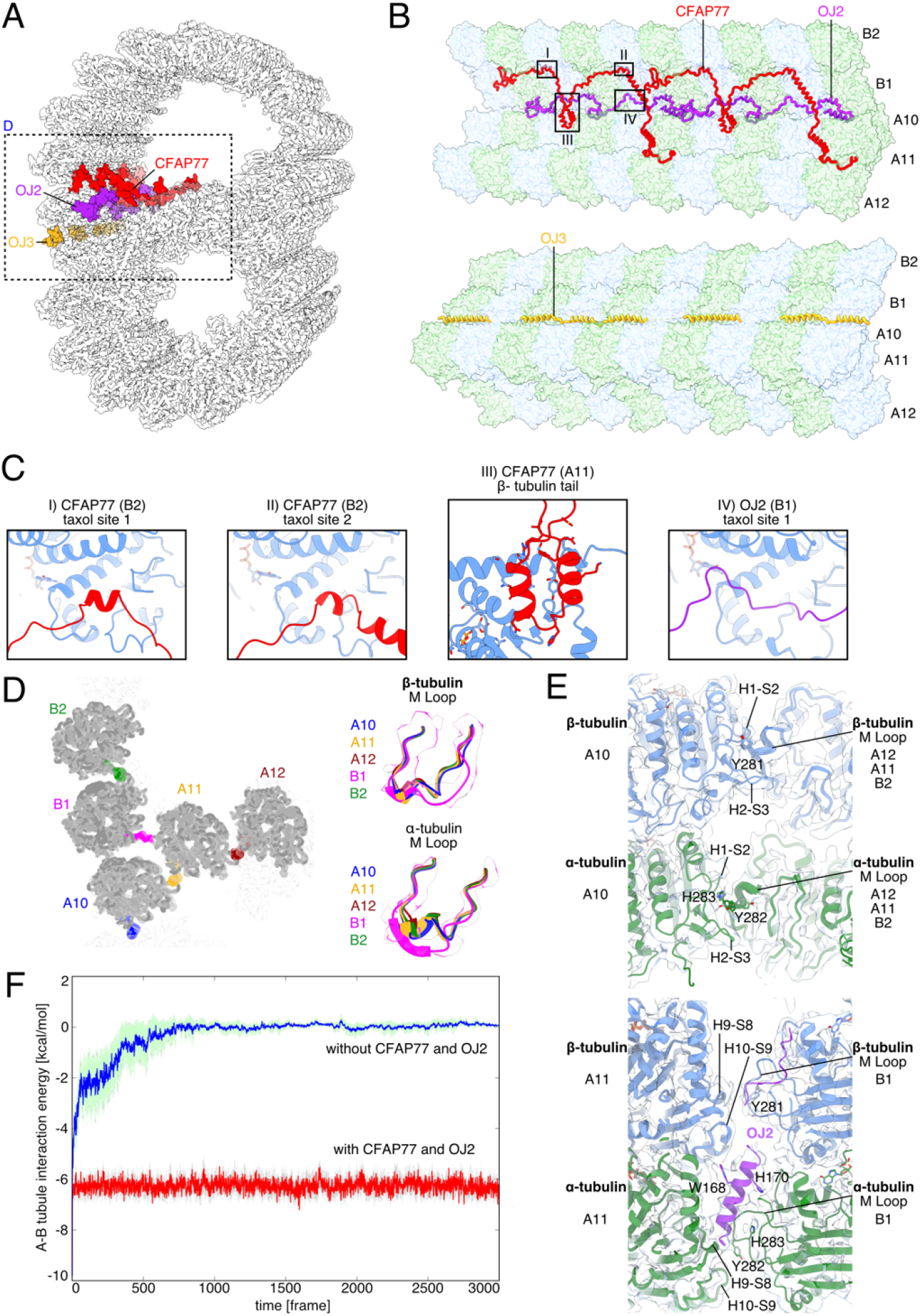
CFAP77 stabilizes the outer junction. (A) Cross-sectional view of the *Tetrahymena* outer junction highlighting proteins CFAP77 (red), OJ2 (purple) and OJ3 (yellow). (B) Architecture of the *Tetrahymena* outer junction including proteins CFAP77, OJ2 and OJ3. (C) Helices of CFAP77 occupy taxane-binding pockets of beta tubulin of PF B2. A loop of OJ2 occupies the taxane-binding pocket of beta tubulin of PF B1. The helix-turn-helix motif of CFAP77 is positioned near the C-terminus of beta tubulin of PF A1. (D) Cross-sectional view of the OJ. M-loops of tubulin are coloured according to protofilament: B2 (green); B1 (purple); A10 (blue); A11 (gold); A12 (dark red). To the right, those same M Loops are superimposed. (E) The canonical (top) and unique (bottom) lateral interactions between tubulin subunits. PFs A11, A12, and B2 are superimposed and adopt the same overall conformation and lateral interactions (top). The lateral interactions between B1 and A11 are particularly unique in *Tetrahymena* because they involve OJ2 (bottom). (F) Molecular dynamics simulations of the outer junction with and without CFAP77 and OJ2.

*Tetrahymena* has two paralogs of CFAP77, CFAP77A and CFAP77B (Supplementary Fig. 5A, Table S2). The mass spectrometry analyses revealed that CFAP77A is almost 3 times more abundant in cilia than CFAP77B. Because of the higher expression level, we decided to model CFAP77A.

The detailed analysis of the outer junction architecture reveals two MIPs, TtCFAP77 and OJ2, interact with four outer junction PFs. TtCFAP77 contacts PFs A10, A11, B1, and B2, while OJ2 contacts PFs A11 and B1 (Fig. 5A, B).

TtCFAP77 and OJ2 are 29 and 20 kDa acidic proteins, respectively, with short alpha helices interspersed with intrinsically disordered regions (Fig. 5B). OJ3 is a filamentous MAP with four alpha helices broken by short intrinsically disordered regions, positioned between PFs A10 and B1. Unlike the 16 nm repeating unit shared by TtCFAP77 and OJ2 spanning two tubulin dimers, OJ3 repeats every 24 nm, spanning three tubulin dimers (Fig. 5B). Each of these three outer junction proteins acts as a thread that weaves across two or more PFs.

Remarkably, two short helices of TtCFAP77 occupy taxane-binding sites on β-tubulin subunits of PF B2 (Fig. 5C). The helix-turn-helix motif of TtCFAP77 is positioned on the β-tubulin C-terminus of PF A11 (Fig. 5C). OJ2 is positioned between PFs B1 and A11 and interacts with the taxane-binding site of a single β-tubulin along PF B1 (Fig. 5C).

We also examine the microtubule binding loop (M-loop) critical for lateral interactions between PFs B1 and A-tubule and their interactions with the outer junction proteins. While the canonical lateral interactions between M-loop and H1-S2 and H2-S3 loops of the adjacent PF (Nogales, Whittaker et al. 1999) are observed throughout the A-tubule and most of the B-tubule, the M-loop of α-tubulin of PF B1 adopts a unique conformation compared to all other M-loops of α-tubulin at the outer junction PFs (Fig. 5D, E). The M-loop of α-tubulin of PF B1 forms hydrogen bonds with both the α-tubulin of A11 and OJ2, providing structural evidence that OJ2 has a stabilising role in the outer junction (Fig. 5E).

To investigate the significance of TtCFAP77 and OJ2 on the outer junction, we used molecular dynamic simulations to examine the behaviour of PF in the presence and absence of TtCFAP77. For the simulation, we used the PFs A10-A12 and B1-B2 as well as TtCFAP77 and OJ2 (see Materials and Methods). Our molecular dynamic simulation showed that tubulins from PFs B1-B2 dissociated from PFs A10-A12 in the absence of both TtCFAP77 and OJ2 (Fig. 5F). Therefore, these MIPs would contribute to the stable binding of the B-tubules to the A-tubule.

The helix-turn-helix motif of TtCFAP77 and the wedge position between PFs A11 and B1 of OJ2 might suppress the intrinsically disordered C-terminal tails of β-tubulin during the assembly of the DMT based on its binding position to PF A11 (Supplementary Fig. 5C). Partial DMTs were previously reconstituted *in vitro* upon the addition of free intact tubulins to subtilisin-treated polymerized microtubules (Schmidt-Cernohorska et al., 2019). Formation of partial DMTs was observed only when microtubules lacked β-tubulin C-terminal tails. The pI of TtCFAP77 and OJ2 are 9.49 and 9.56 respectively, while the pI of β-tubulin is 4.79. Thus, under physiological conditions, these proteins carry positive and negative charges, respectively. It is possible that attractive electrostatic interactions between TtCFAP77, OJ2, and the C-terminal tails of β-tubulin are important for the DMT assembly (Supplementary Fig. 5C). In *Tetrahymena*, OJ2 might contribute more to the suppression of the C-terminal tails of β-tubulin due to its positive surface charge and proximity. The cryo-EM maps of the 48-nm unit of the bovine respiratory cilia and *Chlamydomonas* DMTs have a weak but convincing density for CFAP77 but no equivalent density for OJ2.

### Knockout of CFAP77 moderately reduces *Tetrahymena* cell motility

The structure and molecular dynamic simulation of TtCFAP77 suggested that it plays an important role in the stability of the DMT. Therefore, we investigated how deletion of TtCFAP77 will affect cell swimming, cilia beating, and DMT ultrastructure. Using a germ-line knock-out approach we engineered *Tetrahymena* cells and removed nearly the entire coding region of either *CFAP77A (CFAP77A-KO)* or *CFAP77B (CFAP77B-KO)*, or both genes (*CFAP77A/B-KO*) (Supplementary Fig. 6A, B). Two independent clones were obtained for each mutant. The swimming trajectories of CFAP77 mutants, similar to wild-type cells, are straight but the cell swimming rate is reduced. Single *CFAP77A-KO* and *CFAP77B-KO* mutants covered approximately 66% and 63%, respectively of the wild-type distance while double *CFAP77A/B-KO* mutant travelled 58% of the wild-type distance (Fig. 6A). The analysis of the cilia beating using a high-speed camera revealed that in *CFAP77A/B-KO* mutant ciliary waveform and amplitude were similar to the one observed in the wild-type cells (Fig. 6B) while the ciliary frequency was reduced (Supplementary Fig. 6C, D). Occasionally, in mutants, some asynchronously beating cilia were observed.

**Figure 6.**
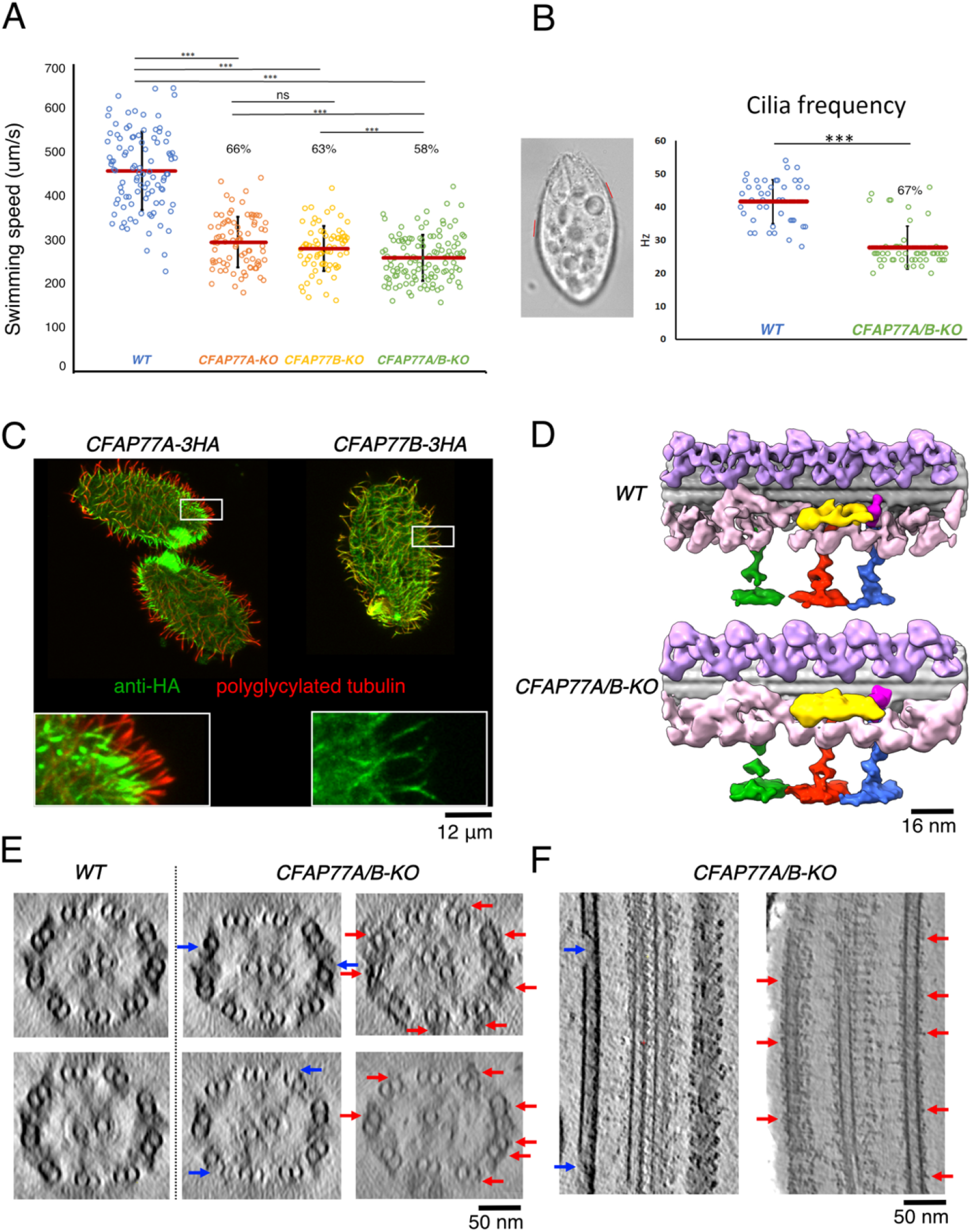
Knockout of *CFAP77A/B* caused mild defects in cilia. (A) Knockout of *CFAP77A* or *CFAP77B*, or both lead up to 40% swimming speed reduction. B) *Tetrahymena* cells with marked exemplary positions (red lines) where cilia beat was analyzed in recorded swimming cells. Graphical representation of measurements of cilia beating frequency in wild-type and *CFAP77A/B-KO* mutants. (C) Differential localization of CFAP77A and CFAP77B in *CFAP77A-3HA* and *CFAP77B-3HA* knock-in mutants. Green - anti-HA; red - poly glycylated tubulin. (D) The 96-nm tomographic maps of *WT (CU-428)* and *CFAP77A/B-KO* showing no abnormalities. (E) The tomographic cross section of WT and *CFAP77A/B-KO* mutants showing the occasional damages in the outer junction of *CFAP77A/B-KO* mutants (blue arrows) and unknown densities near the outer junction region (red arrows). (F) Longitudinal sections from *CFAP77A/B-KO* tomogram showing the outer junction damages (blue arrows) and unknown densities (red arrows).

Immunolocalization of TtCFAP77A and TtCFAP77B proteins expressed as fusions with C-terminal 3HA tags under the control of respective native promoters revealed that TtCFAP77A is present in the proximal end of the cilium while TtCFAP77B localizes along the entire cilium length except for the distal tip (Fig. 6C). However, how TtCFAP77A and TtCFAP77B are targeted for assembly is still unclear.

Using cryo-electron tomography and subtomogram averaging, we obtained structures of the WT and *CFAP77A/B-KO* mutant 96-nm axonemal repeat (Fig. 6D). The averaged tomographic map of the *CFAP77A/B-KO* mutant axoneme looks similar to WT. However, when we examined the raw tomograms of the *CFAP77A/B-KO* mutant, we occasionally observed gaps in outer junction regions (Fig. 6E, F, blue arrows). We also observed more densities binding to the outer junction from outside in the *CFAP77A/B-KO* mutant (Fig. 6E, F, red arrows). Interestingly, the level of tubulin glutamylation in cilia was slightly increased in the *CFAP77A/B-KO* cells compared with the wild-type cells (Supplementary Fig. 7).

## Discussion

In this work, we used cryo-EM to determine the structure of native DMTs from the ciliate *Tetrahymena thermophila* and used a combination of mass spectrometry and artificial intelligence approaches to identify and model proteins inside the DMT. Our structure indicates core MIPs and auxiliary MIPs based on conservation with other species.

### Coordination between the inner and outer surface proteins

Since the DMT was obtained without harsh treatment or enzymatic digestion, its structure represents a close-to-native structure. It shows that there is a difference between periodicity of the outside and inside proteins of the DMT without any clear connection between them. The inside periodicity is a multiple of 16 nm while the outside periodicity is a multiple of 24 nm. In the case of bovine respiratory cilia, it is proposed that Pierce 1 and 2 are the links between the outer 24-nm repeat of the outer dynein arm and the inner 48-nm repeat of MIPs (Gui et al., 2021). Based on the observation of the outer surface filament, we speculate that the outside periodicity can also be achieved independently through a self-regulated head-to-tail arrangement as suggested for the molecular rulers CCDC39 and CCDD40 (Oda, Yanagisawa et al. 2014) and MIPs (Muneyoshi Ichikawa et al., 2019; Ma et al., 2019). Perhaps initial templates of inside and outside proteins at the base of the cilia initiates the perfect registry between the proteins on the inside and outside surfaces.

### Implication of outer surface proteins on intraflagellar transport

The presence of the MAP7-like outer surface filaments in our structure is very interesting since it might have implications of intraflagellar transport on the DMT. As it was shown that MAP7 has biphasic regulation on kinesin-1 due to competitive binding to microtubules (Ferro, Fang et al. 2022), outer surface filaments might impose regulatory effects on dynein and kinesin, partitioning anterograde and retrograde transport tracks. Since there is no evidence that this is a conserved feature in the axoneme (Gui et al., 2021; Ma et al., 2019), this might be a species-specific feature for intraflagellar transport in *Tetrahymena*. On the other hand, a sleeve-like density covering the DMT was observed in the *Chlamydomonas in situ* centriole structure (van den Hoek, Klena et al. 2021), which may prevent the continuous contact of intraflagellar transport motors with the DMT. While the mechanisms of intraflagellar transport to overcome the sleeves in *Chlamydomonas* and the outer surface filaments in *Tetrahymena* DMT are supposed to be different, a fundamental question of how the intraflagellar transport trains can accomplish that feat is an intriguing one.

### The impact of MIPs on ciliary motility

Our study also identifies the conserved protein CFAP77 as an important outer junction protein. CFAP77 is found to localize along the length of human epithelial cilia but is most abundant in the middle (Blackburn, Bustamante-Marin, Yin, Goshe, & Ostrowski, 2017). Expression of human CFAP77 increases during ciliated epithelial cell differentiation (Blackburn et al., 2017). In buffalo sperm, CFAP77 is both phosphorylated and ubiquitinated (P. F. Zhang et al., 2021). Hypomethylation of the CFAP77 gene has also been linked to opioid dependence (Montalvo-Ortiz, Cheng, Kranzler, Zhang, & Gelernter, 2019). It is still unclear how to link post-translation modifications of CFAP77 with its function in cilia.

Molecular dynamic simulation, knockout and mass spectroscopy data in this study suggest that CFAP77 is important for the swimming of *Tetrahymena* probably through its stabilizing role. Interestingly, the phenotype of *CFAP77A/B-KO* is very similar to *RIB72A/B-KO* (Stoddard et al., 2018) and *CFAP115-KO* mutants (Fabritius et al., 2021) with lower ciliary beating frequency and coordination of beating between cilia and excessive curve or kink of some cilia. It is possible that MIPs are required for stabilization of the DMT as the force transducer for proper ciliary beating. Lacking certain MIPs reduces the tensile strength of the DMT, which leads to unexpected curves or kinks during ciliary bending and hence affects the beating coordination. Since the force during ciliary bending is stronger at the base, stability of the axoneme base is likely more important for ciliary function. Coincidently, more TtCFAP77 is present in the proximal half of the cilium (Fig. 6C); possibly the B-tubule is more stable in that region.

### The outer junction assembly

While TtCFAP77 structure and position suggest that it might be important for the suppression of the C-terminal tails of β-tubulins, our study shows that TtCFAP77 is not vital for the assembly of the DMT as the cilia still form with only occasional damages (Fig. 7). The mild phenotype might be because *Tetrahymena* has the species-specific protein OJ2, which might have a redundant role in ciliary assembly or stabilisation. Thus, the phenotype of a CFAP77 KO might be more severe in the case of other species and a potential ciliopathy gene in humans. Based on the increase in polyglutamylation in the *CFAP77A/B-KO* mutants and the presence of proteins outside the DMT, we speculate that the destabilization of the DMT might lead to the increase in recruitment of tubulin tyrosine-like ligases outside (Fig. 7). As the result, the polyglutamylation increase in the DMT might compensate for the destabilization from the lack of CFAP77.

**Figure 7.**
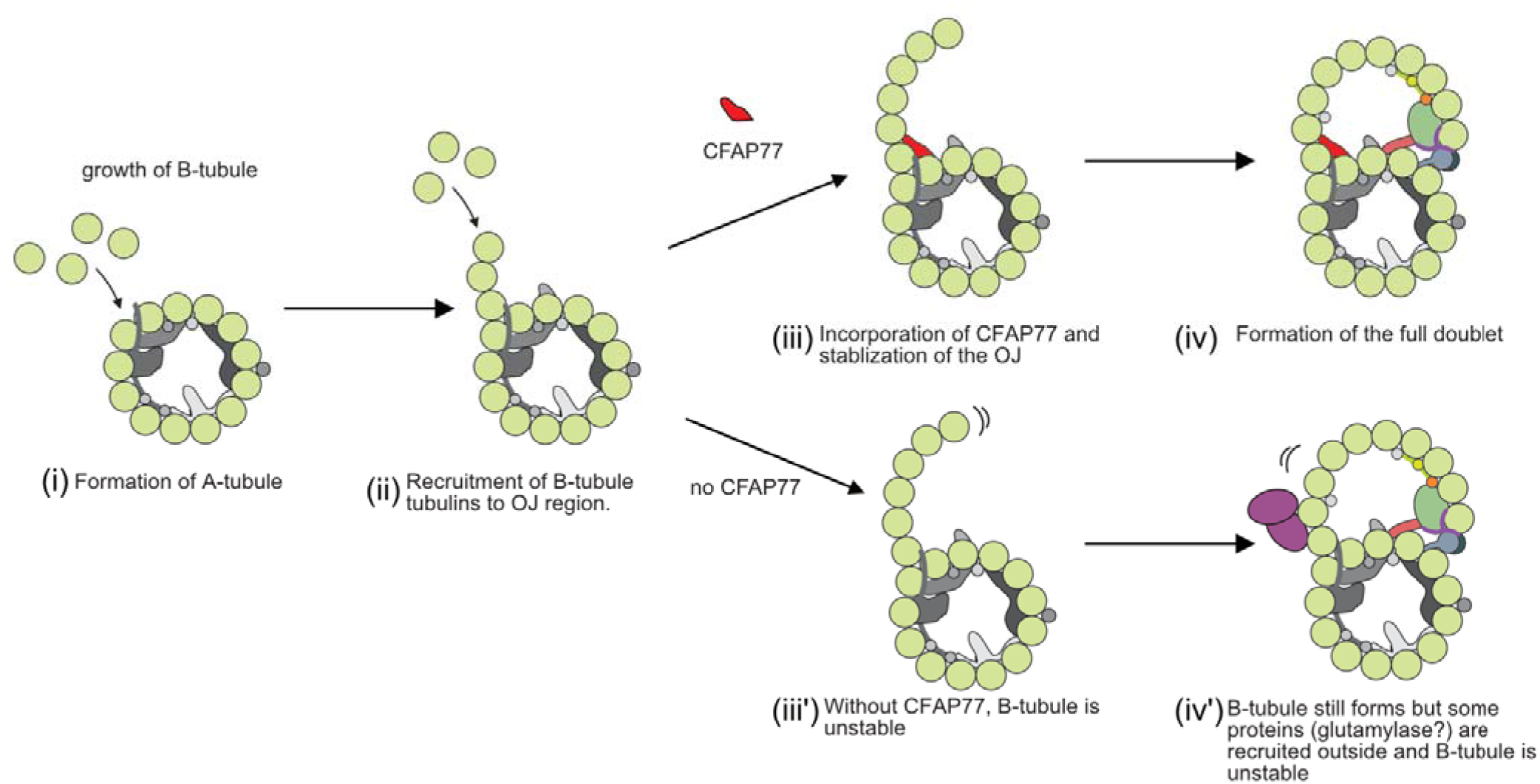
A model of the role of CFAP77 on the formation of the B-tubule. Once A-tubule is formed (i), free tubulins bind to form the hook at the OJ (ii). With CFAP77, the newly form B-tubule hook is stabilized (iii), leading to the final DMT formation (iv). Without the CFAP77, B-tubule is not properly stabilized (iii’) but have additional clusters of proteins on the outer surface (iv’) (likely TTLLs, tubulin tyrosine-like ligases with glutamylase activity) assuming based on the elevated level of tubulin polyglutamylation in cilia.

Interestingly, CFAP77 was not found in the human primary cilia proteome (Mick et al., 2015). Moreover, the orthologous protein is not encoded by the *C. elegans* genome. Yet the DMT exists in the human primary cilia and worm sensory cilia albeit in shorter segments. Therefore, it is likely that in non-motile primary cilia the DMT assembly is CFAP77-independent. It was recently shown that Azami Green-fused Tau-derived peptide added to the microtubules induces formation of the more complex structures, including DMT (Inaba et al., 2022) similar to the doublet reconstitution of C-terminal tail truncated tubulins (Schmidt-Cernohorska et al., 2019). Therefore, it’s possible that other proteins in primary cilia bind to the outer surface of microtubules and interact with or suppress the C-terminal tails of tubulins.

In conclusion, using cryo-EM and improved cilia preparation methods, we have visualized the structure of the native DMT from *Tetrahymena thermophila* and identified DMT positions of 38 MIPs. Moreover, we have shown that an evolutionarily conserved protein, CFAP77, is an outer junction protein and its knockout affects cilia motility (Fig. 7). Our findings contribute to our understanding of MIP function and leave many questions regarding the identity of outer surface proteins and their implication in intraflagellar transport.

## Materials & Methods

### Cell culture

All *Tetrahymena* strains (*CU428* wild-type, *CFAP77A-3HA, CFAP77B-3HA CFAP77A-KO, CFAP77B-KO*, *CFAP77A/B-KO*, *RIB72B-KO*, and *RIB72A/B-KO* mutants) used in this study were grown in SPP media (Williams, Wolfe, & Bleyman, 1980) in a shaker incubator at 30°C and 150 rpm. Cultures were harvested at an OD_600_ of approximately 0.7. Refer to (Black, Dai, Peri, Ichikawa, & Bui, 2021) for further details.

### Cilia isolation using dibucaine treatment

*Tetrahymena* cells were harvested and deciliated as previously described (Black et al., 2021). Briefly, a 2 L cell culture was harvested by centrifugation at 2000 g for 10 minutes at 23°C. Whole cells were resuspended in fresh SPP and adjusted to a total volume of 24 mL. Dibucaine (25 mg in 1 mL SPP) was added, and the culture was gently swirled for 45 seconds. To stop the dibucaine treatment, 75 mL of ice-cold SPP supplemented with 1 mM EGTA was added and the dibucaine-treated cultures were centrifuged for 15 minutes at 2000 g and 4°C. The supernatant (cilia) was collected and centrifuged at 30000 g for 45 minutes at 4°C. The pellet (cilia) was gently washed and resuspended in Cilia Wash Buffer (50 mM HEPES at pH 7.4, 3 mM MgSO4, 0.1 mM EGTA, 1 mM DTT, 250 mM sucrose). The resuspended cilia were flash-frozen with liquid nitrogen and stored at −80°C.

### Purification of the intact axoneme

Intact axonemes were prepared as previously described (Black et al., 2021). The frozen cilia were thawed on ice and then centrifuged for 10 minutes at 8000 g and 4°C. The pellet (cilia) was washed and resuspended in Cilia Final Buffer (50 mM HEPES at pH 7.4, 3 mM MgSO_4_, 0.1 mM EGTA, 1 mM DTT, 0.5% trehalose). NP-40 Alternative (Millipore Sigma #492016) was added to a final concentration of 1.5% for 45 minutes on ice and then the de-membranated axonemes were centrifuged. Intact axonemes (membranes removed with detergent) can be used for tomography or further DMT preparation (see below). For tomography, the pelleted intact axonemes were resuspended in Cilia Final Buffer to a final concentration of 3.8 mg/mL. Crosslinking was done with glutaraldehyde at a final concentration of 0.015%. Bradford reagent (Bio-rad #5000201) was used to measure the total protein concentration.

### Purification of DMTs

DMTs were prepared as previously described (Black et al., 2021; Kubo et al., 2021). The intact axoneme pellet (see above) was resuspended with Cilia Final Buffer and ADP was added to a final concentration of 0.3 mM. The sample was incubated at room temperature for 10 minutes. ATP was added to a final concentration of 1 mM and the sample was again incubated at room temperature for 10 minutes. DMT samples were adjusted to 2.2 mg/mL using Cilia Final Buffer.

### Cryo-EM sample preparation

C-Flat Holey thick carbon grids (Electron Microscopy Services #CFT312-100) were treated with chloroform overnight. For single-particle analysis, DMT sample (4 μl) was applied to the chloroform-treated, negatively glow-discharged (10 mA, 15 s) grids inside the Vitrobot Mk IV (Thermo Fisher) chamber. The sample was incubated on the grid for 15 seconds at 23 °C and 100% humidity then blotted with force 3 for 5 seconds then plunge frozen in liquid ethane.

### Cryo-EM data acquisition

A total of 18,384 movies were collected using the Titan Krios 300 keV FEG electron microscope (Thermo Fisher) equipped with direct electron detector K3 Summit (Gatan, Inc.) and the BioQuantum energy filter (Gatan, Inc.) using SerialEM (Mastronarde, 2005). The movies were collected with a beam shift pattern of four movies per hole and four holes per movement. The final pixel size is 1.370 Å/pixel. Each movie has a total dose of 45 electrons per Å^2^ over 40 frames. The defocus range was between −1.0 and −3.0 μm at an interval of 0.25 μm.

### Image processing

Motion correction and dose-weighting of the movies were performed using MotionCor2 (Zheng et al., 2017) implemented in Relion 3 (Zivanov et al., 2018), and the contrast transfer function parameters were estimated using Gctf (K. Zhang, 2016). Micrographs with apparent drift, ice contamination, and bad contrast transfer function estimation were discarded.

The particles were picked using a combination of manual and automatic pickings to speed up this process. The rare top views of the filaments were picked manually using e2helixboxer (Tang et al., 2007). The side views of the filaments were picked automatically using Topaz by training with a set of side view manual picks (Bepler et al., 2019). The Topaz coordinates were converted into the filament coordinates using clustering and line fitting Python scripts based on RANSAC algorithm.

Particles of 512 × 512 pixels were extracted with 8-nm overlapped, binned twice, and pre-aligned using a modified version of the Iterative Helical Real Space Reconstruction script (Egelman, 2007) in SPIDER (Frank et al., 1996) to work with non-helical symmetry. The alignment parameters were then transferred to Frealign for aligning the particles for six iterations in Frealign (Grigorieff, 2007), and then converted into Relion 3.1. Iterative per-particle-defocus refinement and Bayesian polishing were done for the 8 nm particles.

Each particle was subtracted from its tubulin lattice signal and underwent 3D classification into two classes to obtain the 16-nm repeat particles. The 16-nm repeat particles were then subjected to 3D classification into 3 classes to obtain the 48-nm repeat particles. The 48-nm particles were then refined, resulting in a global resolution of 4.1 Å for WT from 148365 particles respectively.

To improve the local resolution for modelling, we performed focused refinements of different regions by different masks to cover approximately two PFs. Next, the maps were then enhanced by DeepEmhancer (Sanchez-Garcia et al., 2021) to improve visualization and interpretability.

### Modelling

Modelling of tubulins was done as in Yang et al. (Yang et al., 2022) based on PDB 6UOH. For MIP modelling, two strategies were carried out. For conserved MIPs with available models from *Chlamydomonas reinhardtii* (PDB 6U42), homologous models of *Tetrahymena* proteins were either constructed using Modeller (Webb & Sali, 2014) or predicted using ColabFold (Mirdita et al., 2021). The homologous models were fitted into the 48-nm density map of *Tetrahymena* based on the relative location in the *Chlamydomonas* map or by docking using UCSF ChimeraX function fitmap (Pettersen et al., 2021). The final models were then modelled to the density using Coot (Emsley, Lohkamp, Scott, & Cowtan, 2010) and real-space refined in Phenix (Afonine et al., 2018). For unknown densities in the *Tetrahymena* map, connected density was first segmented manually using Chimera. The segmented density was then submitted to DeepTracer server (Pfab et al., 2021) to trace the C-alpha backbone. The C-alpha model was then searched for fold similarity in the library of ColabFold predicted models of all proteins in the proteome of *Tetrahymena* cilia using Pymol *cealign* function. The top candidate ColabFold predicted models were then fitted to the full *Tetrahymena* density map for evaluation. Suitable candidates were gone through modelling in Coot and real-space refinement in Phenix. In certain cases, we used the same density from the *Tetrahymena* map of K40R mutant (Yang et al., 2022) to facilitate modelling since the K40R map has a better resolution at 3.5 Å. All of the maps and model visualization were done using ChimeraX (Pettersen et al., 2021).

### Cryo-ET sample preparation

The axonemes for tomography were cross-linked by glutaraldehyde (final concentration 0.15%) for 40 min on ice and quenched by 1M HEPES. Axonemes were approximately 3.6 mg/mL when mixed with 10 nm gold beads in a 1:1 ratio for a final axoneme concentration of 1.8 mg/mL. Crosslinked axoneme sample (4 μl) was applied to negatively glow-discharged (10 mA, 15 s) C-Flat Holey thick carbon grids (Electron Microscopy Services #CFT312-100) inside the Vitrobot Mk IV (Thermo Fisher) chamber. The sample was incubated on the grid for 15 seconds at 23 °C and 100% humidity then blotted with force 0 for 8 seconds then plunge frozen in liquid ethane.

### Cryo-ET acquisition and reconstruction

Tilt series were collected using the dose-symmetric scheme from −60 to 60 degrees with an increment of 3 degrees. The defocus for each tilt series ranges from −2.5 to −3.5 μm. The total dose for each tilt series is 160 e^-^ per Å^2^. For each view, a movie of 10-13 frames was collected. The pixel size for the tilt series is 2.12 Å. Tomograms were reconstructed using IMOD (Mastronarde, 2005). Frame alignment was done with Alignframes (Mastronarde & Held, 2017). Aligned tilt series were manually inspected for quality and sufficient fiducial presence. Batch reconstruction was performed in IMOD.

### Subtomogram Averaging

Subtomogram averaging of the 4-times binned 96 nm repeating unit of WT and *CFAP77A/B-KO* mutants (2608 and 1702 subtomograms respectively) was done using the “axoneme align” (Bui & Ishikawa, 2013). CTF estimation was done with WARP (Tegunov & Cramer, 2019). Refinement of the 96 nm subtomogram averages was done with the Relion 4.0 pipeline (Kimanius, Dong, Sharov, Nakane, & Scheres, 2021). The resolutions for the 96-nm repeating unit of WT and *CFAP77A/B-KO* DMT are 18 and 21 Å, respectively. For visualization, tomograms were CTF deconvolved and missing wedge corrected using IsoNet (Liu et al., 2021).

### Setting up for CFAP77 coarse-grained MD simulation

We used A10-12 and B1-2 PFs, OJ2 and TtCFAP77 to model *Tetrahymena* Outer Junction (OJ) region. Based on the atomic model of the outer junction region, coarse-grained MD simulation was performed using CafeMol 2.1 (Kenzaki et al., 2011). Each amino acid was represented as a single bead located at its Cα. position in the coarse-grained model. We used the energy functions AICG2+, excluded volume, and electrostatic interaction for predicting dynamics. In the AICG2+, the original reference structure was assumed as the most stable structure, and parameters were modified to represent the interactions in the all-atom reference structure (W. Li, Wang, & Takada, 2014). Three residues (F133, G308, and E401) at the three A10 α-tubulins were anchored for the convenience of the analysis. It is known that the inter α-β dimer interaction is much weaker than intra α-β dimer interaction. Also, inter A-B tubules interaction is weaker than intra A- or B-tubule interaction. To replicate these features in our simulations, we set inter α-β dimer’s interacting force to 0.8 times the original value while the intra α-β dimer’s interacting force was left as the original value (1.0 times). Also, we set inter A-B tubules interacting force to 0.2 times the original value while that of intra A- or B-tubules PFs’ interacting force was set to 0.3 times the original value.

Then, we performed simulations of the outer junction region stability with and without OJ2 and CFAP77 molecules coarse-grained MD, 20 times for each setup. Each MD simulation took 3×10^7 MD steps. Note, that one MD step roughly corresponds to approximately 1ps. The MD simulations were conducted by the underdamped Langevin dynamics at 300K temperature. We set the friction coefficient to 0.02 (CafeMol unit), and default values in CafeMol were used for other parameters.

### CFAP77 gene knock-ins and knock-outs

To reveal the localization of the CFAP77 protein paralogs, we amplified the entire open reading frame with the addition of MluI and BamHI restriction sites at 5’ and 3’ ends, respectively, and 3’UTR with the addition of PstI and XhoI restriction sites at 5’ and 3’ ends, respectively using Phusion HSII High Fidelity Polymerase (Thermo Fisher Scientific Baltics, Lithuania) and primers listed in Table S4, and replaced fragments of the *CFAP44* gene in *CFAP44-3HA* plasmid (Urbanska et al., 2018). Such a construct enabled the expression of the C-terminally 3HA tagged CFAP77 paralogs under the control of their own promoters and selection of the positive clones based on the resistance to paromomycin (neo4 resistance cassette (Mochizuki, 2008)). Approximately 10 μg of plasmids were digested with MluI and XhoI to separate a transgene from the plasmid backbone, precipitated onto DNAdel Gold Carrier Particles (Seashell Technology, La Jolla, CA, USA) according to the manufacturer’s instructions and used to biolistically transform CU428 cells. The positive clones were grown in SPP medium with the addition of an increasing concentration of paromomycin (up to 1 mg/ml) and decreasing concentration of CdCl_2_ (up to 0.2 μg/ml) to promote the transgene assortment.

To knock out *Tetrahymena CFAP77* genes we employed the germline gene disruption approach (Cassidy-Hanley et al., 1997; Dave, Wloga, & Gaertig, 2009). Using primers listed in Table S3 we amplified fragments of the targeted genes with the addition of selected restriction sites and cloned upstream and downstream of the neo4 resistance cassette (Mochizuki 2008) as described elsewhere. In both loci, nearly the entire coding region was removed. To obtain strain with deletion of both *CFAP77A* and *CFAP77B*, we crossed *CFAP77A* and *CFAP77B* heterokaryons and followed with cell maturation and selection as described (Dave et al., 2009). At least two independent clones were obtained for single and double mutants. The loci deletion was confirmed by PCR (primers in Table S3).

### Phenotypic and localization studies

Cells swimming rate and cilia beating were analysed as previously described (Bazan et al., 2021). For protein localization studies, *Tetrahymena* cells were fixed either by adding equal volume of the mixture of 0.5% NP40 and 2% PFA in PHEM buffer or by permeabilization with 0.25 % Triton-X-100 followed by fixation with 4 % PFA (final concentrations) on cover slips, air-dried, blocked using 3%BSA/PBS and incubated overnight with anti-HA antibodies (Cell Signaling Technology, Danvers, MA, USA) diluted 1:200, and polyG antibodies (Duan & Gorovsky, 2002), 1:2000 at 4°C. After washing, 3x 5 min with PBS, samples were incubated with secondary antibodies (anti-mouse IgG conjugated with Alexa-488, and anti-rabbit IgG conjugated with Alexa-555 (Invitrogen, Eugene, OR, USA) both in concentration of 1:300. After washing, the coverslips were mounted in Fluoromount-G (Southern Biotech, Birmingham, AL, USA) and viewed using Zeiss LSM780 (Carl Zeiss Jena, Germany) or Leica TCS SP8 (Leica Microsystems, Wetzlar, Germany) confocal microscope. The expression of the tagged proteins was verified by Western blot as described before (Joachimiak et al., 2021).

### Quantification of Polyglutamylation

Wild-type cells were incubated for 10 min in medium supplied with India Ink to form ink-filled dark food vacuoles. Next, *wild-type* and *CFAP77A/B-KO* cells were fixed side-by-side as described (Wloga et al., 2006) and stained with polyE anti-polyglutamic acid primary antibody (1:2000). Wild-type and mutant cells position next to one another were recorded using confocal microscope and the intensity of the polyE signal in cilia was measured using ImageJ program. For western blotting, 5 μg of cilia protein per lane were separated on a 10% SDS-PAGE and western blots were done as described (Janke et al., 2005), with the primary antibodies at the following dilutions: GT335 anti-glutamylated tubulin mAb (1:1,000), 12G10 anti-α-tubulin mAb (1:40,000), polyE antibodies (1:20 000). The intensity of detected bands was measured using ImageJ and presented as a graph showing a ratio of the glutamylated tubulin to α-tubulin.

### Mass Spectrometry

Samples prepared for cryo-EM were also analysed by mass spectrometry (MS). Approximately 25-30 μg proteins was loaded on the SDS-PAGE gel. Electrophoresis was performed, but the run was terminated before the proteins entered the resolving gel. A band containing all proteins in the sample was then cut out from the gel and subjected to in-gel digestion. Obtained peptides (~2 μg) were chromatographically separated on a Dionex Ultimate 3000 UHPLC. Peptides were loaded onto a Thermo Acclaim Pepmap (Thermo, 75 μm ID × 2 cm with 3 μm C18 beads) precolumn and then onto an Acclaim Pepmap Easyspray (Thermo, 75 μm × 25 cm with 2 μm C18 beads) analytical column and separated with a flow rate of 200 nl/min with a gradient of 2-35% solvent (acetonitrile containing 0.1% formic acid) over 2 hours. Peptides of charge 2+ or higher were recorded using a Thermo Orbitrap Fusion mass spectrometer operating at 120,000 resolutions (FWHM in MS1, 15,000 for MS/MS). The data was searched against *Tetrahymena thermophila* protein dataset from UniProt (https://www.uniprot.org/).

MS data were analyzed by Scaffold_4.8.4 (Proteome Software Inc.). Proteins with mean values of exclusive unique peptide count of 2 or more in the WT MS results were used for analysis. Raw MS data were normalized by total spectra. Student’s *t*-test was applied to K40R and WT MS results using biological triplicates. Proteins exhibiting a minimum of two-fold increase/decrease and a statistical significance threshold (*p* < 0.05) in mutants compared to WT were identified as up- or down-regulated. We used the emPAI calculation of ciliary proteins of specific sizes and periodicities as a filter. We compared the emPAI scores between salt-treated and non-salt-treated axonemes to filter out proteins that are on the surface of the microtubule, rather than the lumen.

## Data Availability

The data produced in this study are available in the following databases:

- Cryo-EM map: EMDB EMD-XXXX
- Model coordinates: PDB XXXX

## Acknowledgements

We thank Drs. Kelly Sears, Mike Strauss, Kaustuv Basu (Facility for Electron Microscopy Research at McGill University) for helping with data collection and Dr. Muneyoshi Ichikawa for critically reading of the manuscript. SK is supported by JSPS Overseas Research Fellowships. KHB is supported by the grants from Canadian Institutes of Health Research (PJT-156354) and Natural Sciences and Engineering Research Council of Canada (RGPIN-2016-04954).

## Additional Information

The authors declare no conflicts of interest.

## Author contributions

Conceptualization: KHB. Methodology: TL, KP, AG, AAZK, CD, MVP, ZF, PH. Formal analysis: SK, CB, EJ, SKY. Investigation: SK, CB, EJ, SKY, TL, KP, AG, AAZK, CD, MVP, ZF, PH. Resources: DW, KHB. Writing–original draft: KHB, DW, SK, CB. Writing–review & editing: KHB, DW, SK, CB, EJ, KP.

## Supplementary Figures

**Supplementary Figure 1.**
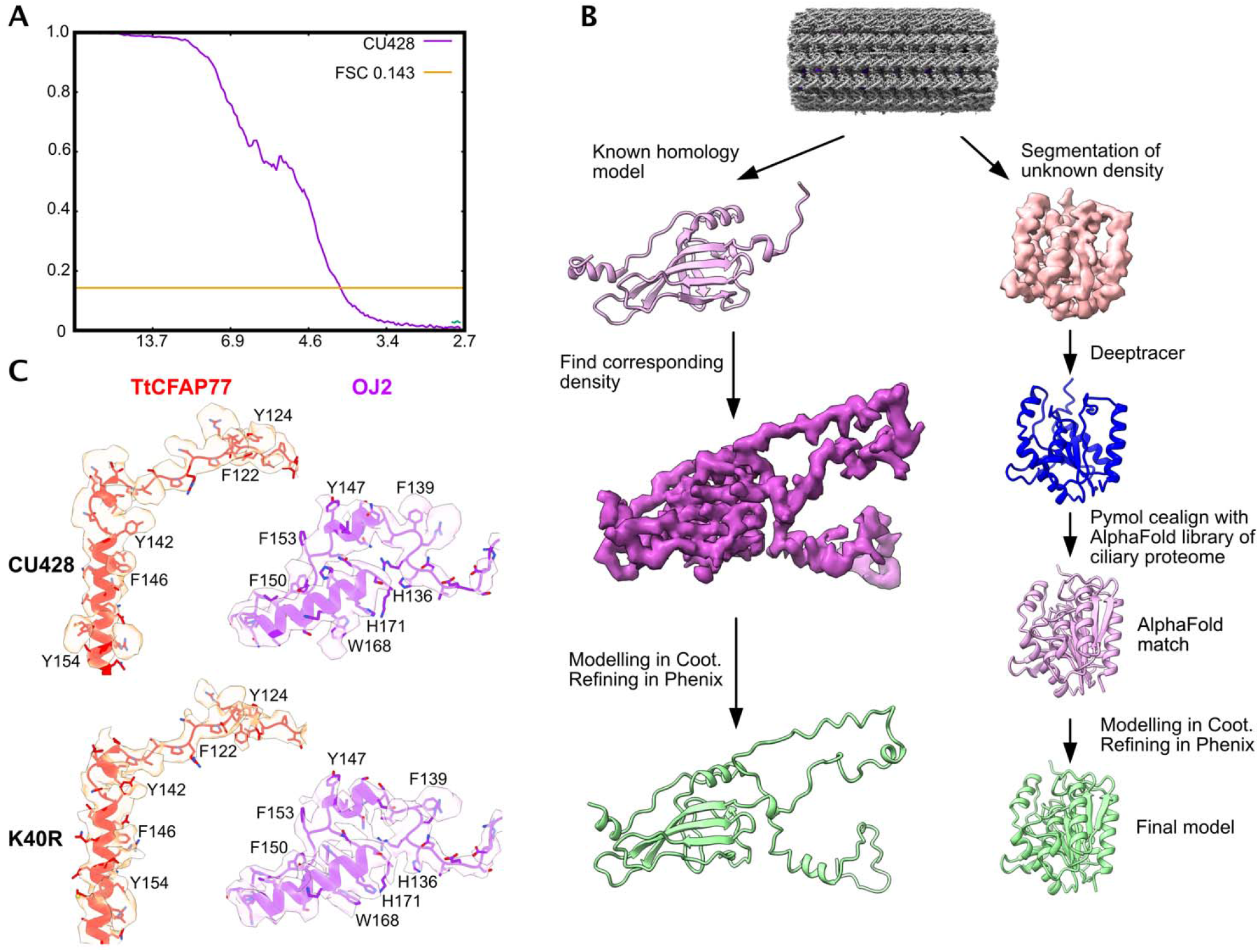
(A) Fourier Shell Correlation of the EM map showing the resolution of 4.1 Angstrom. (B) The modelling workflow to identify and model proteins in our work. (C) Example of the quality of model from CFAP77 and OJ2 in the map of *CU428* (4.1 Angstrom) and α-tubulin K40-acetylation-less mutant, *K40R* (3.7Å) to show the side chain density.

**Supplementary Figure 2.**
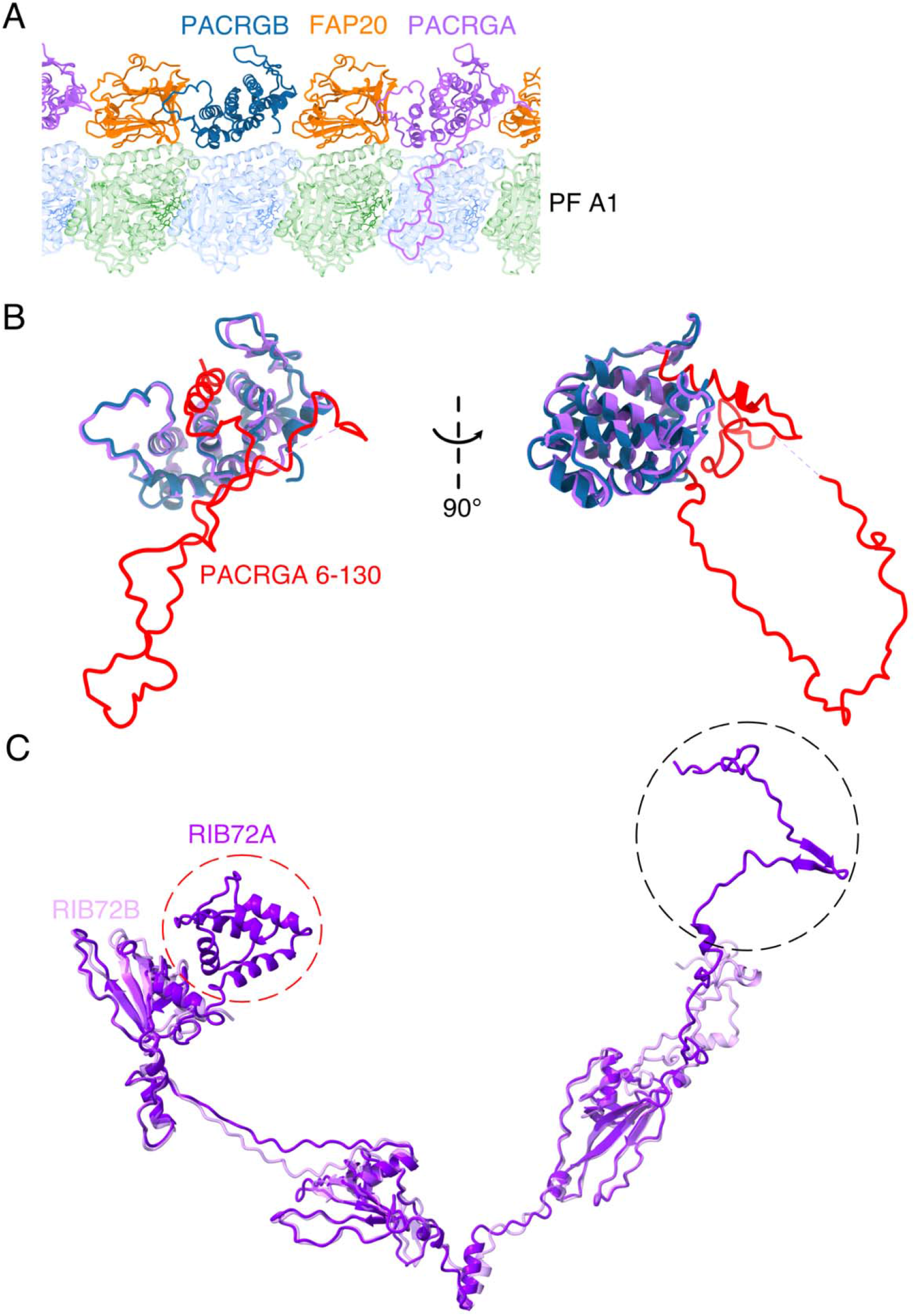
Paralogs of PACRG and RIB72. (A) Arrangement of the IJ filament in *Tetrahymena*. The IJ filament in *Tetrahymena* has a 16-nm periodicity due to the present of PACRGA and PACRGB paralogs. It appears that PACRGA and PACRGB form a dimer with FAP20 but does not form a filament. (B) The difference between PACRGA and PACRGB in the N-terminal domain (highlight in red). (C) The difference between two paralogs RIB72A and RIB72B. The C-terminal, EF-hand domain (black dashed line) and the N-terminal domain (red dashed line) are missing in RIB72B.

**Supplementary Figure 3.**
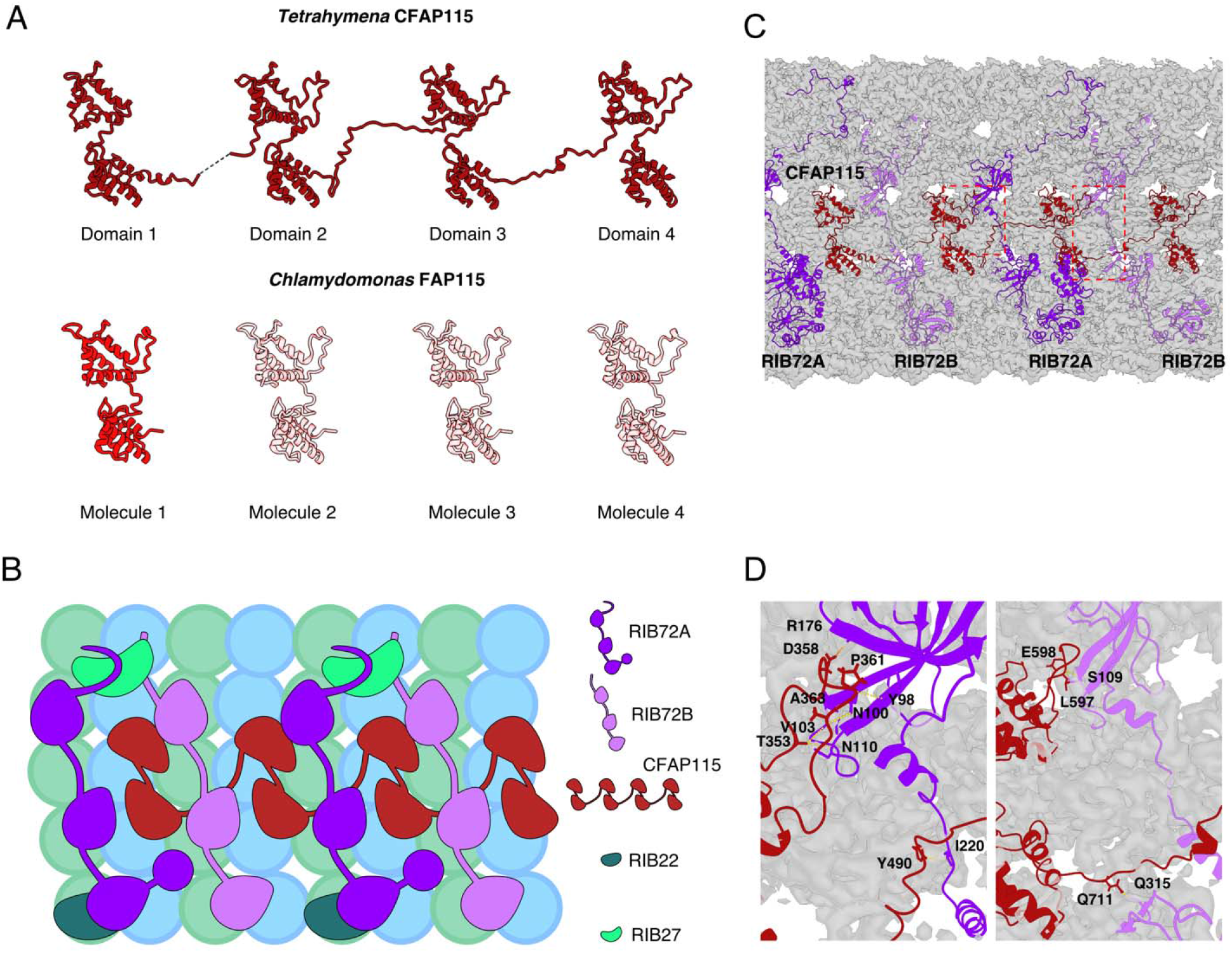
The domain organization of FAP115. (A) Domain organization of TtCFAP115 consisting of 4 tandem EF-hand domain pairs while CrFAP115 has only one EF-hand domain pairs. Thus, TtCFAP115 span 32 nm instead of 8 nm like in the case of CrFAP115. (B) Cartoon depicts the organization of RIB72A, RIB72B, CFAP115, RIB27 and RIB22. Since RIB72A interacts more extensively with RIB22 and RIB27 than RIB72B, RIB22 and RIB27 are only missing in *RIB72AB-KO* mutant but not *RIB72B-KO* mutant. While TtCFAP115 interacts with both TtRIB72A and TtRIB72B, TtCFAP115 is only missing in *RIB72AB-KO* mutant.

**Supplementary Figure 4.**
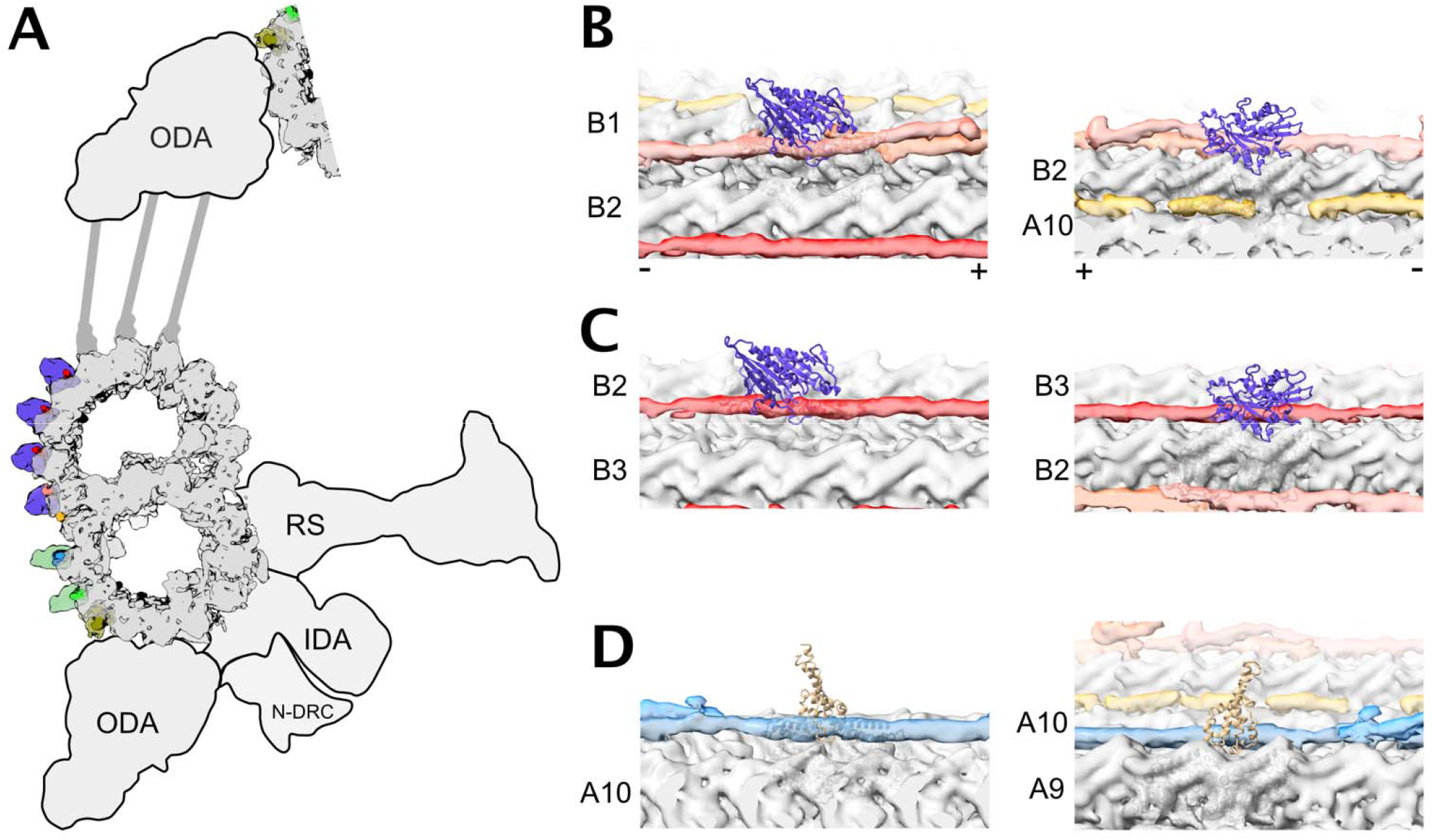
The outer surface filament presents steric clashes with dynein and kinesin. (A) A cartoon scheme of the DMT showing the putative regions for intraflagellar transport between A8-A10 and B1-B5. Violet densities: docking of kinesin microtubule-binding domain onto the B-tubule PFs, green densities: docking of dynein microtubule-binding domain onto the A8 and A9. ODA: Outer dynein arm; RS: Radial spoke; IDA: Inner dynein arm; N-DRC: Nexin-Dynein regulatory complex. (B-D) Docking of kinesin (PDB 6OJQ) on to the outer surface filament B1B2 (B) and B2B3 (C) showing a clear clash. (D) Docking of dynein-2 microtubule-binding domain (PDB 6RZB) onto the filament A9A10 showing the clear steric clash of microtubule-binding domain. (-) and (+) signs indicate the minus and plus ends of the microtubule respectively.

**Supplementary Figure 5.**
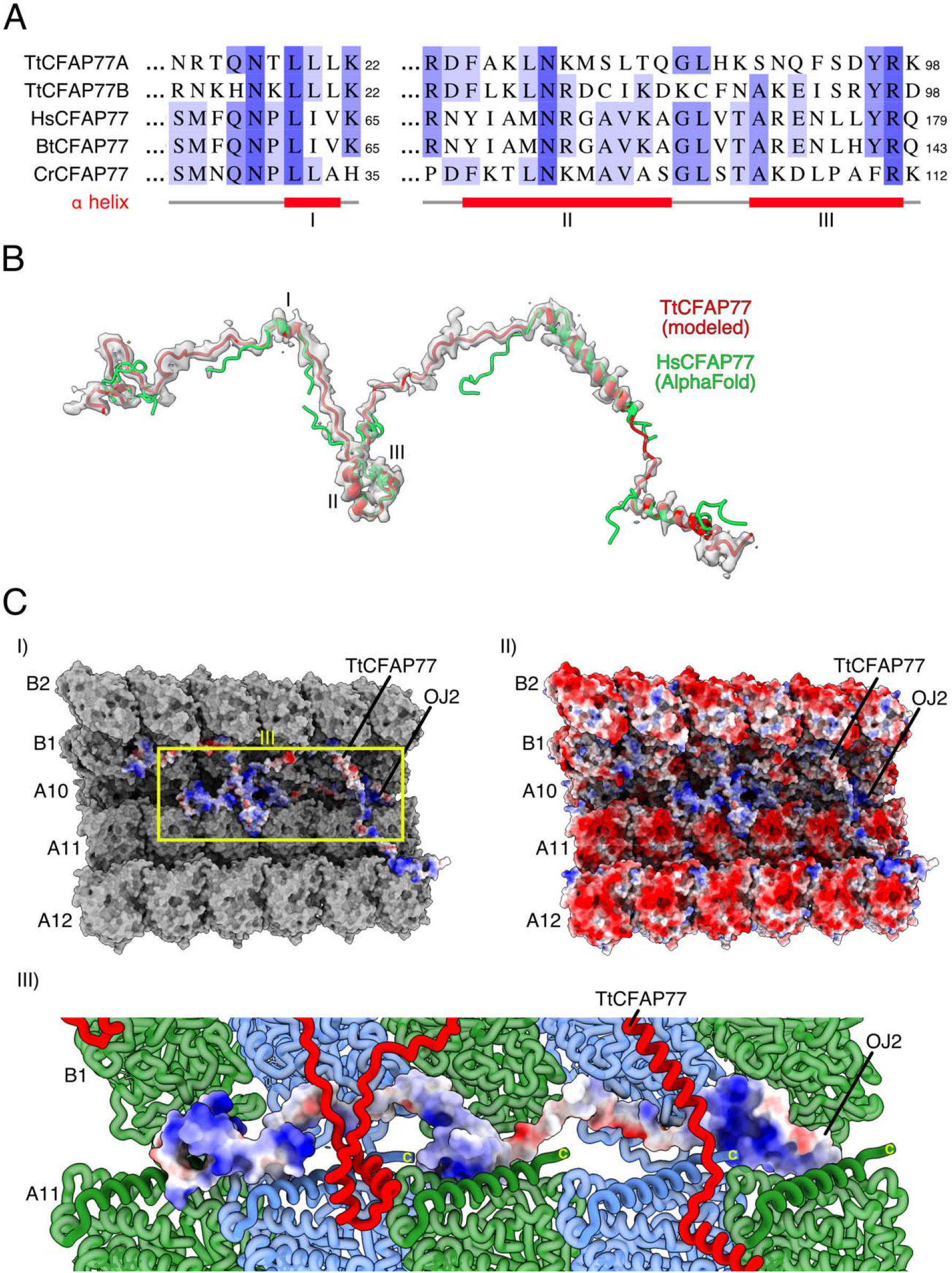
Conservation of CFAP77. (A) Multiple sequence alignment of CFAP77. Human (Hs), bovine (Bt), and *Chlamydomonas* (Cr) sequences as well as two orthologs of *Tetrahymena* (Tt) were included. (B) Cryo-EM model of *Tetrahymena* CFAP77. Folded regions of the AlphaFold model of human CFAP77 (green) were fitted into the map used to model the *Tetrahymena* protein. (C) Panels I and II: Electrostatic potential of the outer junction proteins TtCFAP77 and OJ2. Panel III: Electropositive grooves of OJ2 are proximal to β-tubulin C-termini of PF A11.

**Supplementary Figure 6.**
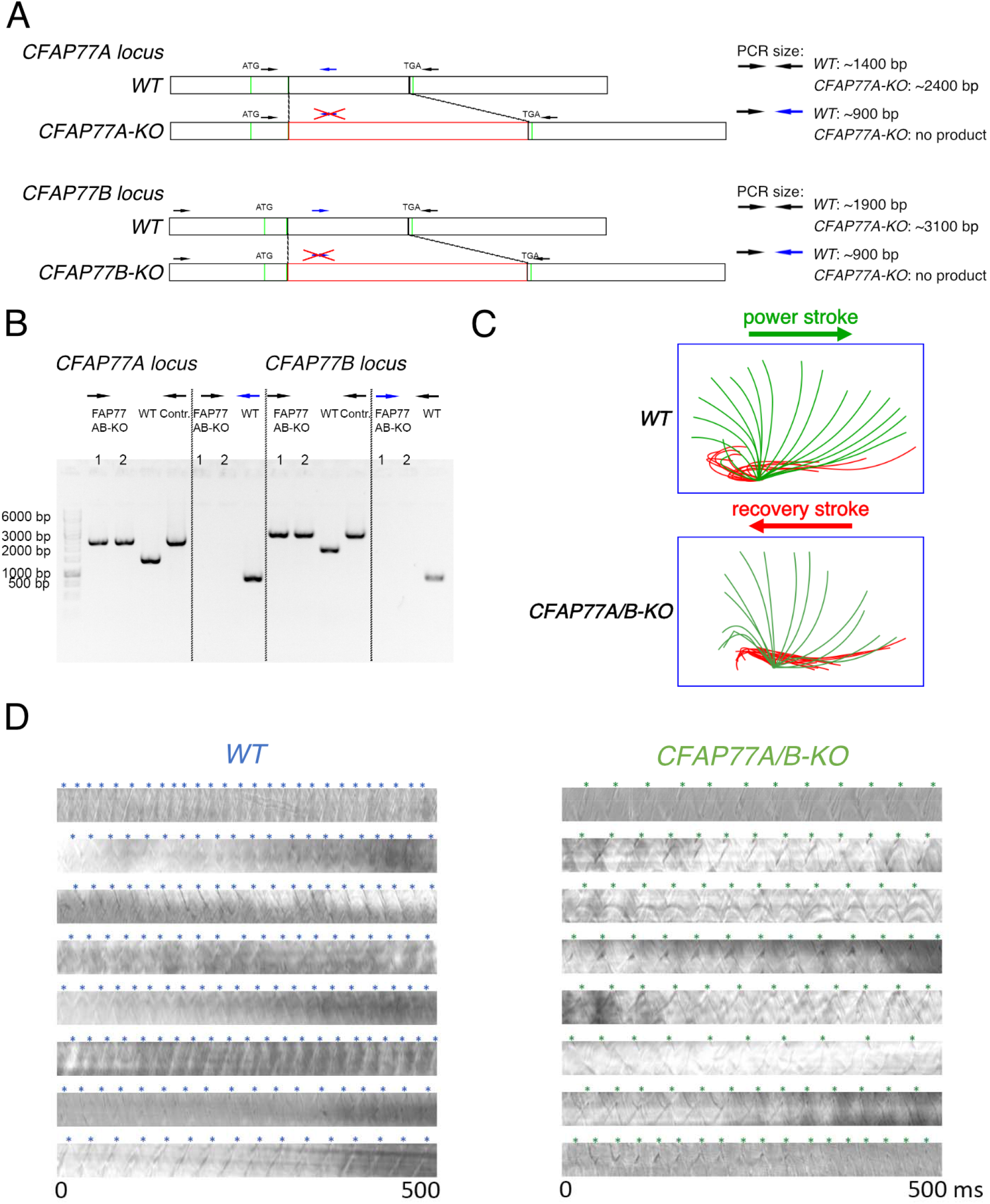
Deletion of two paralogs, CFAP77A and CFAP77B reduces cilia beating frequency. (A) Homologous recombination methodology for *CFAP77A* and *CFAP77B* knockout strains. (B) PCR-based confirmation of the gene knockout. (C) Lack of both CFAP77 paralogs does not apparently alter cilia beating amplitude and waveform. (D) Examples of kymographs of cilia motility, generated from each movie in ImageJ. Each wave peak (*) on a kymograph corresponds to cilia passing through the drawn line.

**Supplementary Figure 7.**
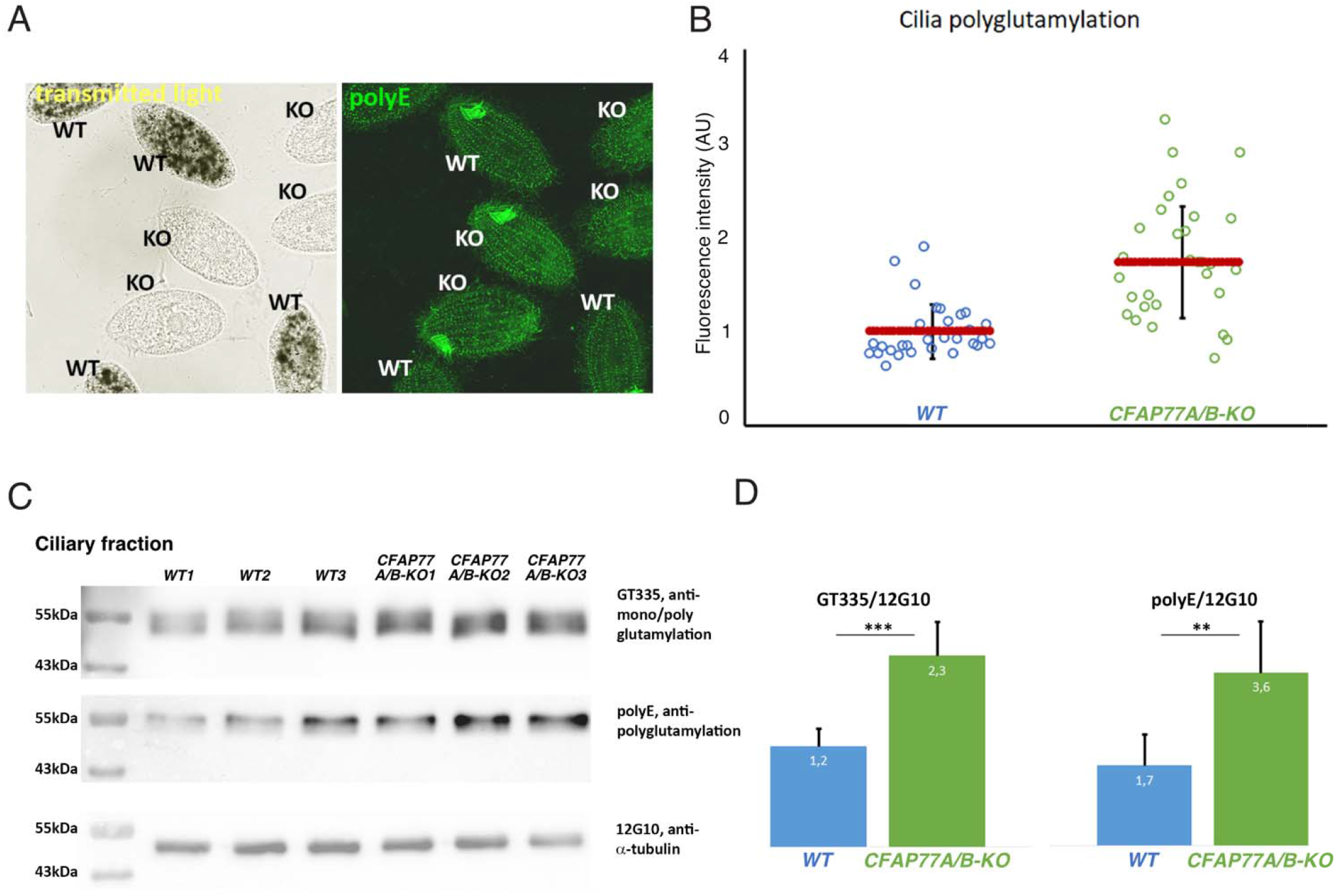
Cilia in the *CFAP77A/B-KO* mutant have a slightly higher level of tubulin glutamylation. Phase-contrast (A, top left) and immunofluorescence (A, bottom left) images of a mixed population of wild-type and *CFAP77A/B-KO* mutant cells stained with polyE antibodies detecting long glutamyl side chains (polyglutamylation). Note that wild-type cells were fed with India Ink and thus contain dark food vacuoles enabling their identification in the population of mixed cells. (B) Graph showing the corresponding quantitative immunofluorescence analyses of the average pixel intensity of axoneme region in mixed and processed side–by–side population of wild-type and mutant cells. (C, D) Western blot (C) and densitometry (graph, D) analyses of the levels of tubulin glutamylation of ciliary tubulin in wild-type and *CFAP77A/B-KO* cells. The level of tubulin was shown using anti-α-tubulin 12G10 antibodies, and levels of tubulin glutamylation were detected using GT335 or polyE antibodies.

**Table S1.** Conserved, non-conserved MIPs in *Tetrahymena thermophila*, vs *Chlamydomonas reinhardtii* vs Bovine. (IJ: Inner junction, OJ: Outer junction, Ribbon: PFs A10-A13). (*: not model but only model the other paralog, **: not model but homolog identified, and similar density found at the same place)

|  | Common MIPs |  |  | Species specific |  |  |
| --- | --- | --- | --- | --- | --- | --- |
| Region | *Tetrahymena* | *Chlamydomonas* | Bovine | Tetra-specific | Chlamy-specific | Bovine-specific |
| A-tubule | TtCFAP21A*<br>TtCFAP21B | FAP21 | CFAP21 | TtB3B4 | FAP68 | EFCAB6 |
| A-tubule | TtCFAP53 | FAP53 | CFAP53 | TtB4B5 | FAP85 | TEKTIN 1 |
| A-tubule | TtCFAP67 | FAP67 | NME7 | TtB5B6 | FAP90 | TEKTIN 2 |
| A-tubule | TtCFAP115 | FAP115 |  | TtIJ34 | FAP166 | TEKTIN 3 |
| A-tubule | TtCFAP127 | FAP127 | MNS1 | TtMIP21A | FAP182 | TEKTIN 4 |
| A-tubule | TtCFAP129** | FAP129 |  | TtMIP25A | FAP222 | TEKTIP1 |
| A-tubule | TtCFAP182 | FAP182 | PIERCE1 | TtMIP26A | FAP252 |  |
| A-tubule |  |  | PIERCE2 | TtOJ2 | RIB21 |  |
| A-tubule | FAM166B | - | FAM166B | TtRIB22 | RIB30 |  |
| B-tubule | TtCFAP45 | FAP45 | CFAP45 | TtRIB26 | FAP68 |  |
| B-tubule | TtCFAP112 | FAP112 |  | TtRIB27 | FAP85 |  |
| B-tubule | TtCFAP210 | FAP210 | CFAP210 | TtRIB35 | FAP166 |  |
| IJ | TtCFAP20 | FAP20 | CFAP20 | TtRIB38 | FAP182 |  |
| IJ | TtCFAP52 | FAP52 | CFAP52 | TtRIB57 | FAP222 |  |
| IJ | - | FAP126 | FLTP |  | FAP252 |  |
| IJ | - | FAP276 | CFAP276 |  | FAP273 |  |
| IJ | TtPACRGA | PACRG | PACRG |  | FAP363 |  |
| IJ | TtPACRGB |  |  |  | RIB21 |  |
| OJ | TtCFAP77A | FAP77 | CFAP77 |  | RIB30 |  |
|  | TtCFAP77B |  |  |  |  |  |
| OJ | TtOJ3 | - | FAM183A |  |  |  |
| Ribbon | - | FAP95 | CFAP95 |  |  |  |
| Ribbon | TtCFAP106A* | FAP106 | ENKUR |  |  |  |
| Ribbon | TtCFAP106B |  |  |  |  |  |
| Ribbon |  | FAP107 | CFAP107 |  |  |  |
| Ribbon | TtCFAP141 | FAP141 | CFAP141 |  |  |  |
| Ribbon | TtCFAP143 | FAP143 | SPAG8 |  |  |  |
| Ribbon | TtCFAP161A | FAP161 | CFAP161 |  |  |  |
|  | TtCFAP161B* |  |  |  |  |  |
| Ribbon | TtRIB43a_S | RIB43a | RIBC2 |  |  |  |
| Ribbon | TtRIB43a_L |  |  |  |  |  |
| Ribbon | TtRIB72A | RIB72 | EFHC1 |  |  |  |
| Ribbon | TtRIB72B |  | EFHC2 |  |  |  |

**Table S2.** UniprotID and Gene name of the MIPs in this study.

| **MIP** | **UniprotID** | **Gene name** |
| --- | --- | --- |
| TtRIB72A | I7M0S7 | TTHERM_00143690 |
| TtRIB72B | I7MCU1 | TTHERM_00584850 |
| TtCFAP115 | Q23KF9 | TTHERM_00193760 |
| TtCFAP161A | Q22WJ6 | TTHERM_00155380 |
| TtCFAP161B | I7M8Z8 | TTHERM_00219350 |
| TtCFAP182 | Q24BV4 | TTHERM_01049330 |
| TtRIB43A_S | A4VDZ5 | TTHERM_00641119 |
| TtRIB43A_L | Q240R7 | TTHERM_00624660 |
| TtCFAP67 | W7XGD1 | TTHERM_000372529 |
| TtCFAP21A | I7MK20 | TTHERM_00340080 |
| TtCFAP21B | I7MLS4 | TTHERM_00494280 |
| TtCFAP53 | Q23YQ8 | TTHERM_01207630 |
| TtCFAP143 | A4VD56 | TTHERM_00502359 |
| TtCFAP52 | Q22ZH2 | TTHERM_01094880 |
| TtCFAP45 | W7XCX2 | TTHERM_001164064 |
| TtCFAP210 | Q23EX8 | TTHERM_00643530 |
| TtCFAP112 | Q23A15 | TTHERM_00884620 |
| TtPACRGA | I7MLV6 | TTHERM_00446290 |
| TtPACRGB | I7M317 | TTHERM_00499570 |
| TtCFAP20 | Q22NU3 | TTHERM_00418580 |
| TtIJ34 | I7M9T0 | TTHERM_00348390 |
| TtCFAP141 | A4VCU8 | TTHERM_00242188 |
| TtCFAP106A | I7M279 | TTHERM_00137550 |
| TtFAP106B | I7LU20 | TTHERM_00370750 |
| TtFAP127 | I7LV70 | TTHERM_00338110 |
| TtOJ3 |  |  |
| TtCFAP77A | Q22WR6 | TTHERM_00974270 |
| TtCFAP77B | Q239Q2 | TTHERM_01190430 |
| TtOJ2 | Q236L2 | TTHERM_00086850 |
| TtRIB27 | I7LUL4 | TTHERM_00522810 |
| TtRIB57 | I7ME23 | TTHERM_00406590 |
| TtRIB35 | I7ME81 | TTHERM_00267880 |
| TtRIB26 | Q232I6 | TTHERM_00594110 |
| TtRIB22 | I7LT67 | TTHERM_00691900 |
| TtRIB38 | Q23JL9 | TTHERM_00758860 |
| FAM166B | Q235M9 | TTHERM_01052990 |
| TtMIP26A | Q238X3 | TTHERM_00449470 |
| TtMIP25A | Q22B75 | TTHERM_01109800 |
| TtMIP21A | Q237T1 | TTHERM_00320040 |

**Table S3.** Mass spectrometry analysis of wild type (WT), *RIB72B* and *RIB72A/B* knockout mutants showing the missing proteins. (Only proteins with quantitative value > 1 are shown).

| UniprotID | *WT (CU428)* |  |  | *RIB72B-KO* |  |  | *RIB72A/B-KO* |  |  |
| --- | --- | --- | --- | --- | --- | --- | --- | --- | --- |
|  | S1 | S2 | S3 | S1 | S2 | S3 | S1 | S2 | S3 |
| I7MCU1 (**RIB72B**) | 217.4 | 199.6 | 212.8 | 0.0 | 0.0 | 0.0 | 0.0 | 0.0 | 0.0 |
| I7M0S7 (**RIB72A**) | 172.9 | 189.0 | 181.7 | 246.0 | 245.9 | 242.0 | 0.0 | 0.0 | 0.0 |
| Q23KF9 (**FAP115**) | 300.3 | 313.5 | 297.0 | 294.5 | 281.9 | 281.9 | 0.0 | 0.0 | 0.0 |
| Q238X3 (**MIP26A**) | 101.3 | 102.4 | 115.2 | 84.5 | 89.0 | 102.7 | 0.0 | 0.0 | 0.0 |
| Q235M9 (**FAM166B**) | 35.8 | 34.4 | 44.3 | 33.8 | 39.2 | 38.8 | 0.0 | 0.0 | 0.0 |
| I7LT67 (**RIB22**) | 31.4 | 30.9 | 37.2 | 16.9 | 21.2 | 21.7 | 0.0 | 0.0 | 0.0 |
| I7LUL4 (**RIB27**) | 28.8 | 20.3 | 31.9 | 13.7 | 27.6 | 30.8 | 0.0 | 0.0 | 0.0 |
| Q22CT6 | 27.9 | 36.2 | 39.0 | 1.1 | 4.2 | 3.4 | 0.0 | 0.0 | 0.0 |
| W7XC77 | 13.1 | 11.5 | 16.0 | 7.4 | 8.5 | 12.6 | 0.0 | 0.0 | 0.0 |
| I7LV80 | 10.5 | 12.4 | 9.8 | 0.0 | 0.0 | 1.1 | 0.0 | 0.0 | 0.0 |
| Q22TY0 | 9.6 | 8.8 | 8.9 | 0.0 | 0.0 | 0.0 | 0.0 | 0.0 | 0.0 |
| I7LVP2 | 8.7 | 11.5 | 9.8 | 0.0 | 1.1 | 2.3 | 0.0 | 0.0 | 0.0 |
| Q231F9 | 5.2 | 1.8 | 7.1 | 1.1 | 0.0 | 1.1 | 0.0 | 0.0 | 0.0 |
| Q22CT4 | 5.2 | 4.4 | 3.5 | 0.0 | 0.0 | 1.1 | 0.0 | 0.0 | 0.0 |
| I7M2F8 | 2.6 | 4.4 | 2.7 | 0.0 | 41.3 | 3.4 | 0.0 | 0.0 | 0.0 |
| I7M6E1 | 2.6 | 3.5 | 1.8 | 0.0 | 0.0 | 0.0 | 0.0 | 0.0 | 0.0 |
| Q22KN2 | 1.7 | 2.6 | 1.8 | 0.0 | 0.0 | 0.0 | 0.0 | 0.0 | 0.0 |
| Q22HI8 | 1.7 | 0.9 | 0.9 | 0.0 | 0.0 | 0.0 | 0.0 | 0.0 | 0.0 |
| I7MDS6 | 1.7 | 0.9 | 1.8 | 0.0 | 0.0 | 0.0 | 0.0 | 0.0 | 0.0 |
| I7LWU5 | 1.7 | 0.9 | 1.8 | 1.1 | 1.1 | 0.0 | 0.0 | 0.0 | 0.0 |
| Q22S92 | 1.7 | 0.0 | 0.9 | 0.0 | 0.0 | 0.0 | 0.0 | 0.0 | 0.0 |

**Table S4.**
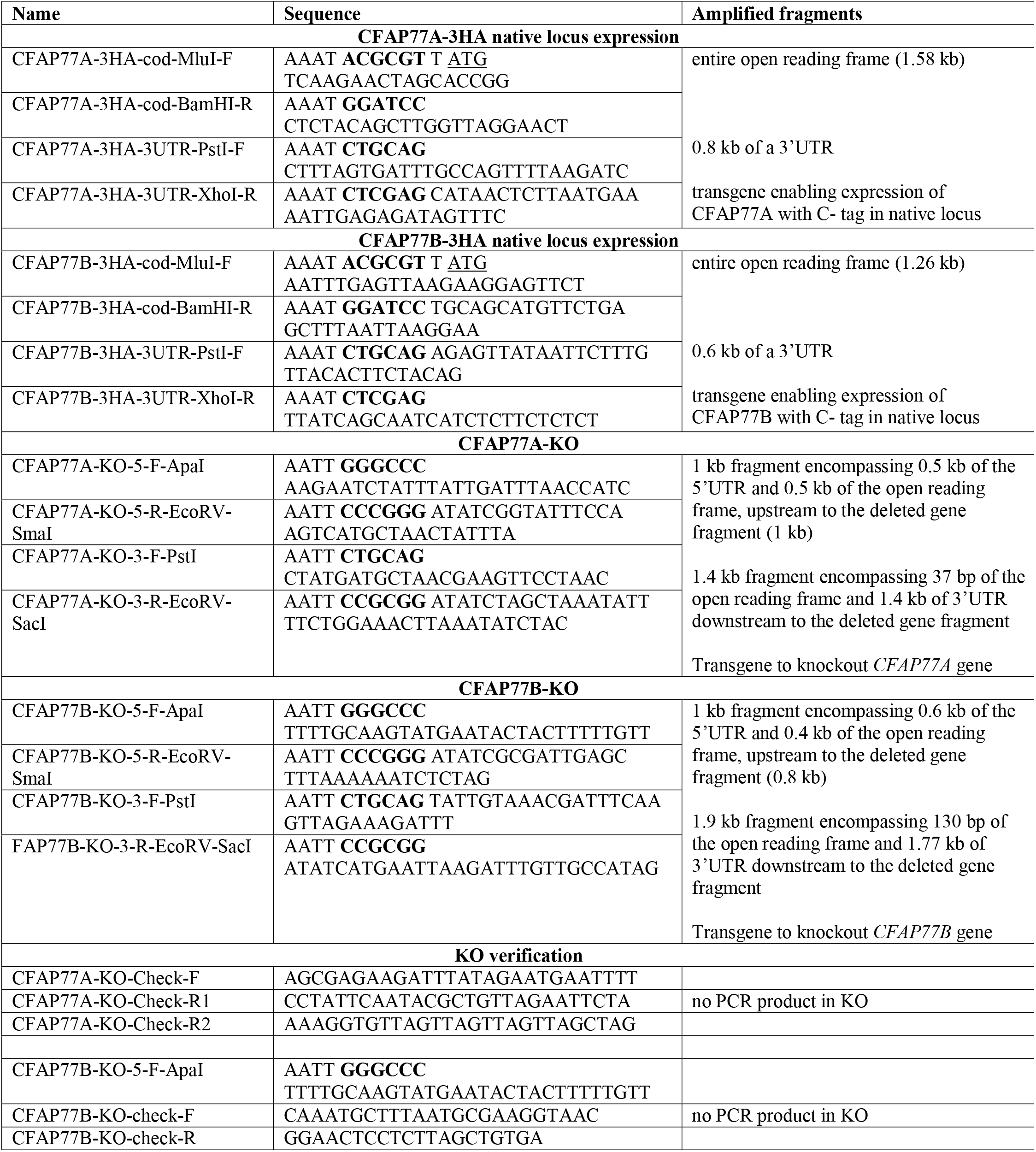
Primers used to engineer *Tetrahymena CFAP77* paralogs mutant cells. Sequence recognized by restriction endonuclease in bold, ATG or TGA are underlined.

## Supplementary Movies

**Supplementary Movie 1:** Highlighting the structure of the native DMT from *Tetrahymena* cilia.

**Supplementary Movie 2:** An overview of the outer junction of the DMT form *Tetrahymena* cilia and their interactions.

